# pH Gradient Drives Redox Reactions in the Nucleolar Caps to Modulate Nucleolar Integrity and Function

**DOI:** 10.64898/2026.11.14.766641

**Authors:** Suibin Ma, Xiang Zhan, Fan Wu, Zhen Mao, Junlin Chen, Jierui Guo, Wenbin Li, Chuan-Qi Zhong, Yongyou Zhang, Xin Zhang, Jiajia Guo, Bo Wang

## Abstract

While enzymatic catalysis is the main driver of redox processes in cells, biomolecular condensates exhibit unique electrochemical properties that can potentiate redox reactions. Nevertheless, the regulatory mechanisms governing the electrochemical equilibrium of biomolecular condensates and their influence on the emergent physicochemical functions remain poorly understood. Here, we demonstrate that specific nucleolar stressors induce alkaline nucleolar caps, which generate an interfacial pH gradient and electrical potential that drive spontaneous redox reactions. Non-enzymatic reactive oxygen species (ROS) production depends on positively charged constituents within the nucleolar caps and correlates strongly with the pH microenvironment. Excessive ROS directly modulates the partition and diffusion of proteins rich in redox-sensitive residues, whereas scavenging ROS markedly accelerates the recovery of nucleolar multiphase organization and functionality upon stress removal. Collectively, our findings propose a non-enzymatic redox mechanism operating within cellular condensates and highlight the critical role of physical interfaces in orchestrating the chemical dynamics of biomolecular condensates.

## Introduction

Biological systems depend on finely orchestrated chemical reactions to sustain cellular functions and adapt to environmental fluctuations. Within this complex network, redox reactions occupy a pivotal role in energy conversion, cellular signaling, and the regulation of metabolic pathways^1^. Conventionally, enzymatic catalysis has been regarded as the primary driver of redox processes in living cells. However, accumulating evidence now supports the existence of non-enzymatic redox mechanisms that operate within both abiological coacervates and biological assemblies^2–13^. These reactions are frequently modulated by physical interfaces, which generate distinct electrochemical microenvironments—ultimately reshaping the energy landscape of biomolecular interactions and redirecting reaction pathways^2–13^.

Biomolecular condensates, formed via phase separation, exhibit unique physicochemical properties that set them apart from their surrounding cellular milieu. Notably, condensates in specialized regions such as the nucleolus display localized variations in pH and ionic composition^14–16^. Recent studies have further elucidated that the net charge of nucleolar proteins plays a critical role in establishing a pH gradient across distinct nucleolar layers^14^. This interfacial pH difference drives the selective distribution of ions within the condensate’s dense phase and facilitates the formation of an electric double layer (EDL)^9^. The EDL, commonly found and best described at liquid– solid or liquid–air interfaces^11,17^, serves as a compensatory mechanism to offset ionic gradients, thereby achieving electrochemical potential equilibrium. These electrochemical features of the condensate microenvironment imply that biomolecular assemblies are not merely structural compartments but chemically active entities capable of modulating chemical functions. One of the most striking consequences of this activity is the potentiation of redox reactions, driven by the EDL-dependent interfacial electric field inherent to condensates^5,7–9,11^. This electric field lowers the energy barrier for chemical reactions-including redox processes-without relying on enzymatic catalysts^13^. Thus, the non-enzymatic redox modulation provides a novel conceptual framework for understanding how cells spatiotemporally regulate biochemical reactions, expanding our perspective beyond traditional enzymatic control mechanisms.

However, while these discoveries reveal the potential of electrochemical environments to influence biochemical dynamics and function, the exact mechanisms by which biomolecular condensates regulate these environments, especially in complex, biologically relevant conditions, remain largely unknown. In this study, we explore how specific nucleolar stressors induce the formation of alkaline nucleolar caps, the microenvironment of which is crucial for restoring the structural and functional integrity of the nucleolus. The formation of these alkaline caps drives spontaneous redox reactions, via generating an interfacial electrochemical potential. This non-enzymatic reactive oxygen species (ROS) generation is reliant on charged constituents within the nucleolar caps and correlates strongly with the pH microenvironment. The excessive ROS in turn regulates the partition and diffusion of proteins enriched in cysteine and methionine Importantly, scavenging ROS accelerates the restoration of nucleolar multiphase architecture and functionality. These observations provide a novel non-enzymatic mechanism for redox reactions in complex heterogeneous biological assemblies, offering new insights into how electrochemical environments within condensates regulate chemical reactions and cellular homeostasis.

## Results

### DNCs potentiate redox reactions and produces ROS by establishing interfacial pH gradient

The nucleolus normally adopts a multiphase architecture, but can undergo nucleolar segregation under various stressed conditions, including those that disrupt rRNA transcription^18–21^. In addition, many proteins, most of which are implicated in mRNA splicing, transiently assemble into a subcompartment known as the dark nucleolar caps (DNCs) that is in close contact with the segregated nucleolus^18^. Our prior data suggest that the DNC exhibits an alkaline microenvironment^21^. Based on recently developed theories^5,12^, the pH difference (ΔpH) between the DNCs and the surrounding milieu is predicted to trigger selective ion distribution within the DNC and the establishment of an electric double layer (EDL) to offset the resulting ionic gradient to achieve phase equilibrium, which is characterized by the equilibrium of the electrochemical potential. By lowering the energy barrier, the interfacial electric field conferred by the existing EDL between the DNC and the surrounding milieu potentiates chemical reactions, including redox reactions. We therefore hypothesized that DNCs enable non-enzymatic redox reactions by establishing an interfacial pH gradient (**Figure 1A**).

**Figure 1.**
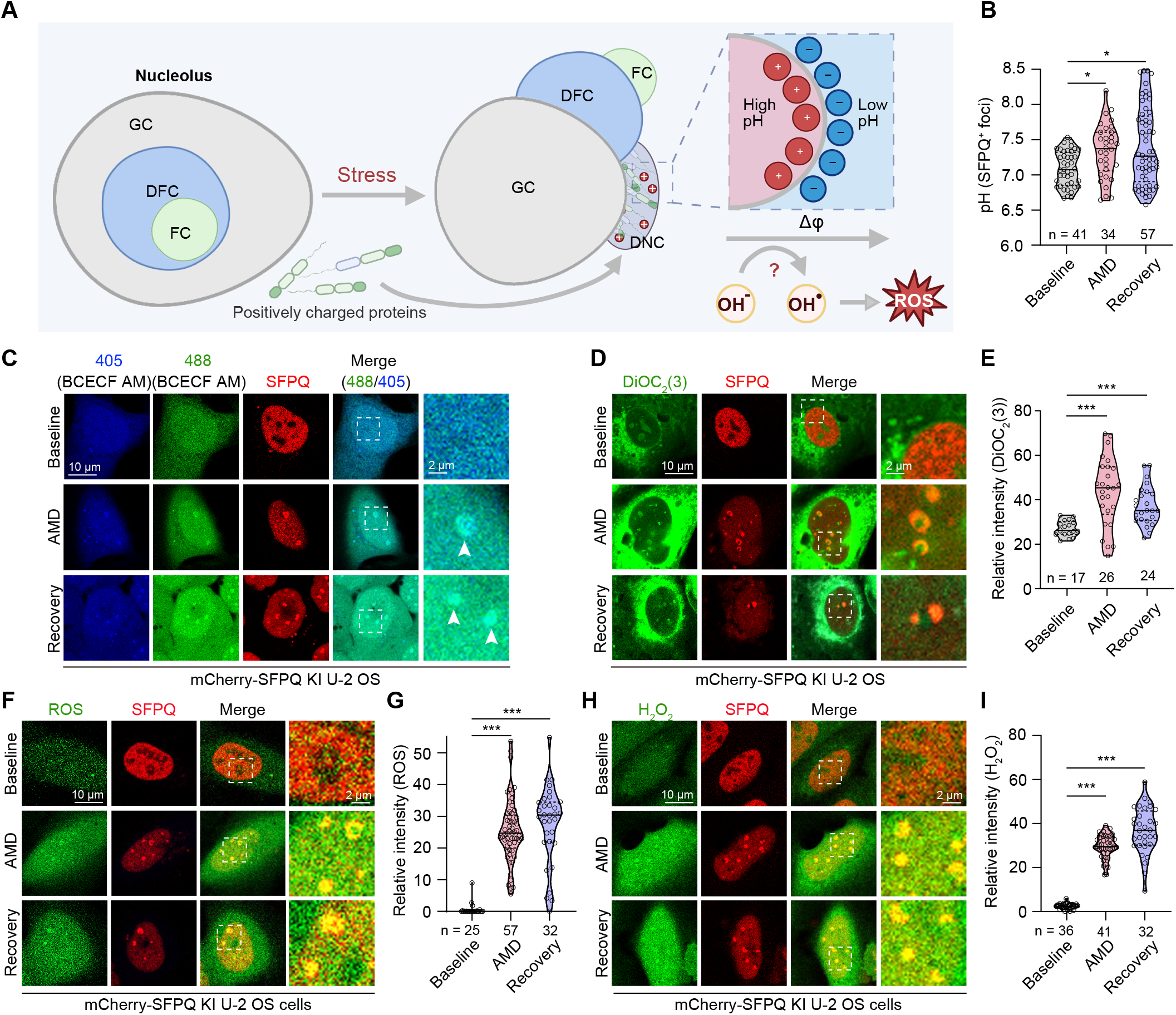
DNCs enable redox reactions and produces ROS by establishing interfacial pH gradient. (**A**) Schematic illustration of the central hypothesis of this study. (**B**, **C**) Live cell imaging of mCherry-SFPQ KI U-2 OS being treated with AMD and recovered in normal growth condition with a pH indicator BCECF-AM. pH was quantified by calculating the ratio of the fluorescence intensity of 488/405 nm in the DNC marked by mCherry-SFPQ (**B**). **(D, E)** Live-cell imaging was performed on mCherry-SFPQ KI U-2 OS cells following AMD treatment and subsequent recovery under normal growth conditions using a potential responsive dye DiOC_2_(3). The fluorescence intensity of DiOC_2_(3) in the DNCs (marked by mCherry-SFPQ) was quantified (**E**). (**F**, **G**) Live-cell imaging was performed on mCherry-SFPQ KI U-2 OS cells following AMD treatment and subsequent recovery under normal growth conditions, utilizing the ROS-sensitive probe CM-H_2_DCFDA. The fluorescence intensity of ROS within mCherry-SFPQ-labeled DNCs was quantified in (**G**). (**H**, **I**) Live-cell imaging of mCherry-SFPQ KI U-2 OS cells after AMD treatment and recovery under normal growth conditions, using the specific H_2_O_2_ fluorescent probe pentafluorobenzenesulfonyl fluorescein. H_2_O_2_ levels was quantified by measuring fluorescence intensity at 488 nm (**I**). For (**B**), (**E**), (**G**), and (**I**), statistical significance was determined using one-way ANOVA (Kruskal-Wallis test). ns, not significant; *p < 0.05; **p < 0.01; ***p < 0.001.

We first sought to delineate the pH dynamics within the DNC microenvironment. We performed live cell imaging to quantitatively measure the DNC pH and the associated nucleolar structure by utilizing a membrane-permeable dye BCECF-AM^22,23^. This fluorescence dye shows a pH-sensitive emission when it is excited at a wavelength of ∼490 nm and a pH-insensitive emission when excited at a wavelength of ∼440 nm (**Figure S1A**). We were able to draw a linear progression between the pH values of standard buffers and the ratiometric intensity of 488/405 nm (**Figure S1B**). Therefore, pH can be quantitatively determined by calculating the emission ratio of wavelengths 488 and 405 nm and fitting it into the standard curve. Given that pH varies within the nucleolar sub-phases^14,16^, we tagged the endogenous loci of *SFPQ*, *NPM1*, *Fibrillarin/FBL*, and *TCOF*, which encoded proteins residing in the DNCs and nucleolar GC, DFC, and FC, respectively, with the mCherry coding sequence in U-2 OS cells^21^, and used the mCherry fluorescence as a mask to directly quantify the stress-dependent pH fluctuation. We ensured that the DNCs (SFPQ^+^) represented a distinct subcompartment from GC, DFC, and FC (**Figure S1C**), as was described previously^18^. Of note, the DNCs were robustly alkalinized not only by AMD treatment (**Figures 1B, 1C**) but also by several other stressors that impeded rRNA biogenesis, including UV and doxorubicin exposure (**Figures S1D-S1G**), whereas no significant alkalinization within the NMP1^+^ GC, FBL^+^ DFC or TCOF^+^ FC subcompartments was detected (**Figures S1H-S1M**). Note that the vast majority of the measured pH values of the DNCs and nucleolar subcompartments were within the calibrated range (5.80-8.73).

As described above, the pH gradient between the DNC and the surrounding microenvironment will elicit an enrichment of specific ion species in the DNC and the emergence of an EDL at the surface of the DNCs^4^, which generates a surface potential, also known as the zeta potential^24^. Currently, the most widely used approach to detect electrical potential involves the use of chemical probes that change their electronic structure in response to changes in the local electric field, which results in a change in their fluorescence property and ratiometric fluorescence^25,26^. Using one of such fluorescent dyes, DiOC_2_(3), we observed that the DNC electrical potential was significantly higher than the surrounding nucleoplasm (**Figures 1D and 1E**). We confirmed that this dye was a reliable indicator of the electrical potential as its detection of the mitochondrial membrane potential was severely abolished by CCCP, a decoupler of the mitochondrial proton gradient (**Figures S1N and S1O**). We also chose to perform an alternative, distinct measurement that can potentially reflect the electric field changes in the DNCs. Micropolarity was recently proposed as a determinant for the structural organization of biomolecular condensates^27^. We hypothesized that the alkaline microenvironment of the DNCs and the presence of a charged surface would reduce its micropolarity (that is, more hydrophobic)^28^. To test this hypothesis, we utilized a 7-sulfonamide benzoxadiazole fluorophore probe, SBD-Halo^27^, whose fluorescence lifetime negatively correlates with polarity. Using a standard curve generated *in vitro*, the micropolarity can be semi-quantitatively measured by fluorescence lifetime imaging microscopy (FLIM). The micropolarity of the DNCs was correlated to a dielectric constant value of approximately 15 using two independent DNC marker proteins, SFPQ and SRSF1 (**Figures S1P and S1Q**). In contrast, the translocated FC region (TCOF^+^) was much more hydrophilic (**Figures S1P and S1Q**). These results were consistent with the observation that the DNC region was more basic than FC under this experimental condition.

To determine if the DNC electrical potential potentiates spontaneous redox reaction, leading to the generation of ROS. We employed a reduction-sensitive probe, CM-H_2_DCFDA, whose fluorescence intensity increases upon exposure to ROS (**Figure S2A**). Interestingly, AMD exposure resulted in prominent ROS generation, which was mostly detected in the DNCs (**Figures 1F and 1G**). Note that the ROS intensity was higher at the recovery stage than that of the AMD treatment, likely due to the continuous generation of ROS during recovery. To validate the results, we used roGFP1, a ratiometric genetic reporter of ROS^29^, which also indicated significant ROS accumulation caused by AMD treatment, as evidenced by the significantly increased intensity ratio of 405 nm to 488 nm (**Figures S2B and S2C**). Importantly, even a lower dose of AMD treatment (0.1 µM) resulted in the generation of ROS in the DNCs (**Figures S2D and S2E**). We further observed the ROS was not distributed in the NPM1^+^ GC, FBL^+^ DFC or TCOF^+^ FC subcompartment (**Figures S2F-S2K**). Together, these data imply that the production of ROS is positively correlated with the pH difference across DNC interfaces. Other stressors that create an interfacial pH gradient in the DNCs, including UV and doxorubicin exposure, similarly caused the specific formation of ROS in the DNCs (**Figures S2L and S2M**). We also confirmed that this interfacial pH gradient-dependent generation of ROS was independently detected in several cell types, such as HeLa, 293T and HCT 116 cells (**Figures S2N-S2S**), suggesting that this is not a cell type-specific phenomenon.

Studies on the ubiquitous electrochemical activity at the liquid-air and liquid-solid interfaces show that an electrical field can transform hydroxide ion (OH^-^) dissociated from a water molecule into a hydroxyl radical (OH•) and a solvated electron^11,17^. The recombination reaction between hydroxyl radicals can then generate H_2_O ^11,17^. Therefore, we utilized a selective probe, pentafluorobenzenesulfonyl fluorescein (PBSF), whose fluorescence intensity increases upon exposure to H_2_O_2_. Indeed, significantly higher levels of H_2_O_2_ were detected in the DNCs (**Figures 1H and 1I**).

### Uncovering the chemical mechanisms of ROS generation in the DNCs

To define the chemical identity of the DNC ROS, we first used ROS scavengers, including N-acetylcysteine (NAC) and glutathione (GSH), which were sufficient to eradicate the ROS (**Figures 2A, 2B, S3A, and S3B**). Further, the selective expression of Catalase, which was fused in frame with the SRSF1 RRM domain for DNC targeting^21^, largely abolished the fluorescent signals in the DNCs (**Figures 2C and 2D**), confirming the chemical identity of ROS. To elucidate the mechanism of H_2_O_2_ generation, we treated the cells with a hydroxyl radical scavenger, L-carnosine, given that the generation of hydroxyl radical (OH•) is a defining intermediate reactant of H_2_O_2_ production driven by the interfacial electrical fields. The addition of L-carnosine significantly diminished the generation of ROS in the DNCs (**Figures 2E and 2F**), suggesting the involvement of hydroxyl radical. To ask if metal-dependent redox chemistry (e.g., Fenton-type reactions) contributes to the ROS in the DNCs, we expressed the metal-chelating protein MT1A. Interestingly, MT1A overexpression in the DNCs did not appreciably reduce the ROS levels (**Figures S3C and S3D**). Together, our data suggest an electrical field-driven, non-enzymatic synthesis of ROS, which is sharply distinct from the conventional reaction mechanisms of ROS in membrane-bound organelles such as mitochondria.

**Figure 2.**
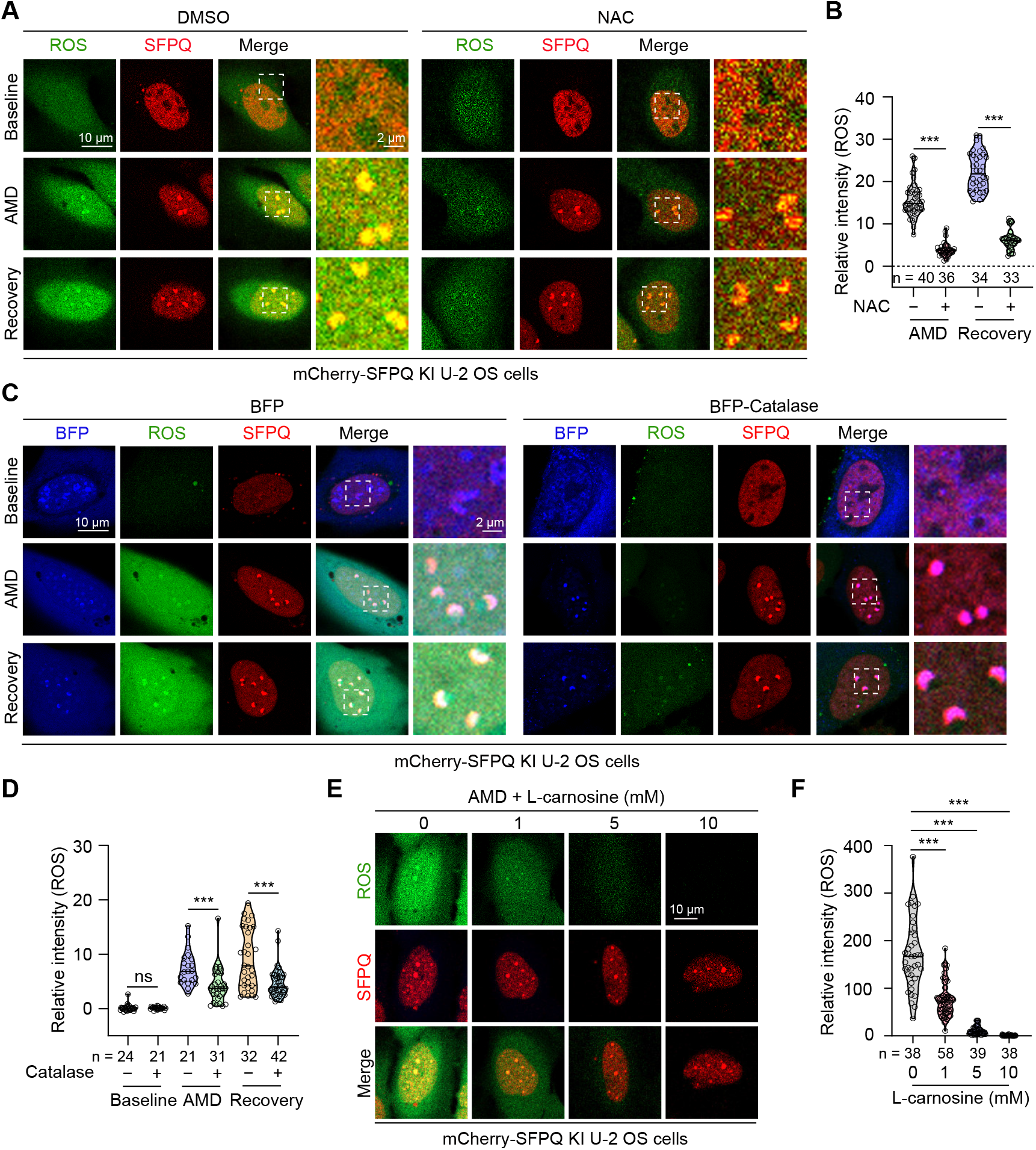
Uncovering the chemical mechanisms of ROS generation in the DNCs. (**A, B**) Live-cell imaging of mCherry-SFPQ KI U-2 OS cells treated with AMD in the presence of DMSO or NAC, and subsequently recovered under normal growth conditions. The fluorescence intensity of ROS was quantified (**B**). (**C, D**) Live-cell imaging was performed on mCherry-SFPQ KI U-2 OS cells expressing BFP-catalase following AMD treatment and subsequent recovery under normal growth conditions. The fluorescence intensity of ROS was quantified (**D**). (**E, F**) mCherry-SFPQ KI U-2 OS cells were cotreated with AMD and a concentration gradient of a hydroxyl radical scavenger, L-carnosine, (0, 1, 5, 10 mM), and the ROS intensity within mCherry-SFPQ-labeled DNCs was assessed via live cell imaging with CM-H_2_DCFDA (**F**). For (**B**) and (**D**), statistical significance was determined using a two-tailed unpaired Student’s *t*-test. For (**F**), statistical significance was determined using one-way ANOVA. ns, not significant; *p < 0.05; **p < 0.01; ***p < 0.001.

### DNC proteome is enriched with positively charged residues

To dissect the physicochemical basis of the electrical potential across the DNC interfaces, we aimed to delineate the biochemical characteristics of the DNC proteome by means of Turbo-mediated proximity labeling. We generated fusion proteins of Turbo and SFPQ or SRSF1, two RNA-binding proteins with high DNC partition^18^. Upon inducing the DNC formation, biotinylated proteins are purified by streptavidin beads and analyzed by label-free quantitative mass spectrometry (**Figure 3A**). We validated that both SFPQ-Turbo and SRSF1-Turbo showed DNC localization only when cells were stressed with AMD, and that biotinylation was specifically observed only when Turbo was engineered (**Figures S4A-S4D**).

**Figure 3.**
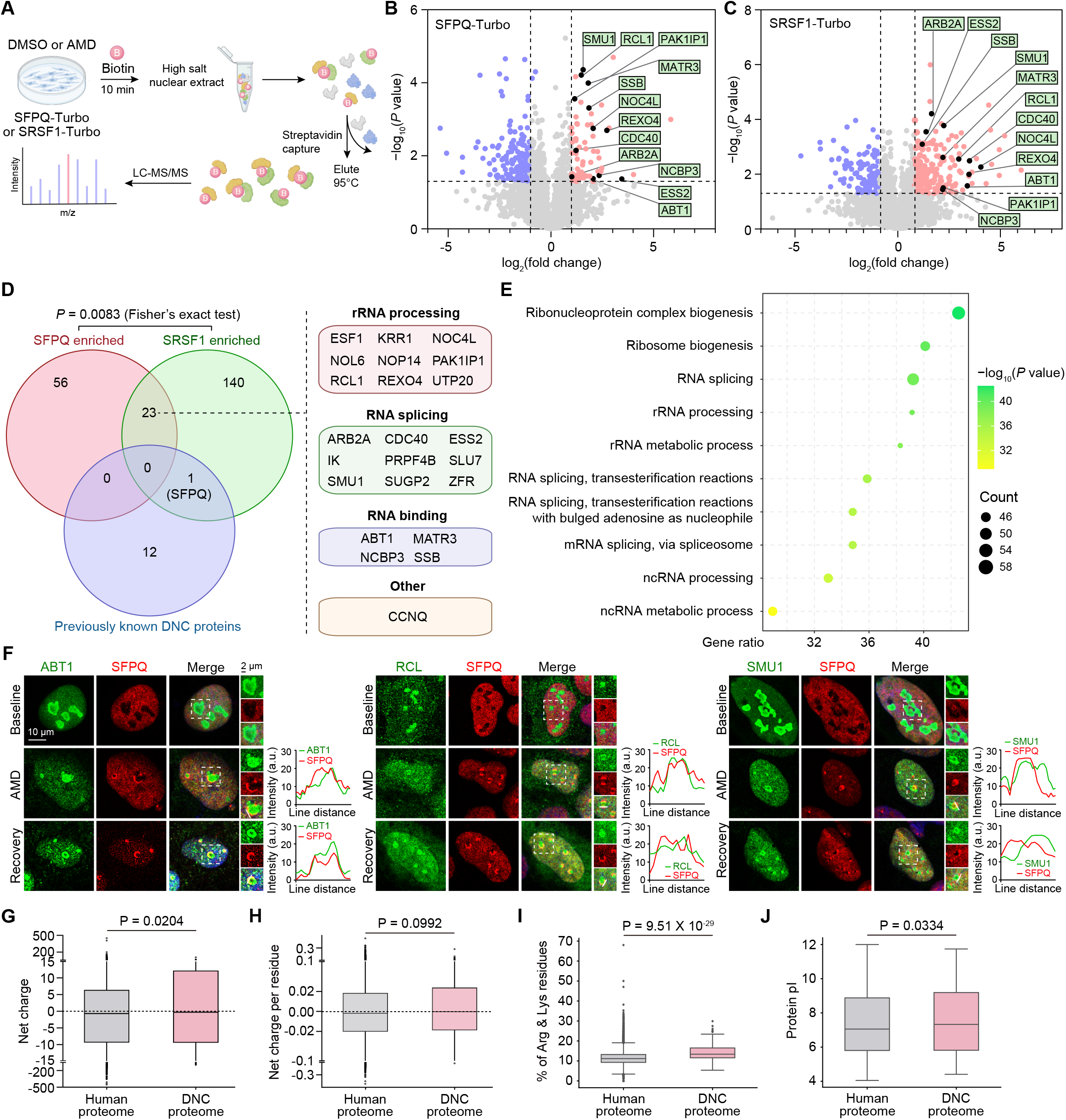
DNC proteome is enriched with positively charged residues. (**A**) Schematic illustration of the streptavidin–mass spectrometry (MS) experimental design for SFPQ-Turbo or SRSF1-Turbo. (**B**, **C**) Volcano plots displaying proteins enriched in MS experiments for SFPQ-Turbo (**B**) or SRSF1-Turbo (**C**). Vertical dashed lines indicate a 2-fold change threshold; horizontal dashed lines denote a P value of 0.05. (**D**) Venn diagram comparing proteins enriched in SFPQ-Turbo and SRSF1-Turbo experiments, and previously known DNC protein. The 23 commonly enriched proteins implicated in rRNA processing, RNA splicing, and RNA binding are listed. (**E**) Gene ontology (GO) analysis of proteins enriched in the MS experiments. (**F**) DNC localization of the indicated proteins was validated by immunofluorescence. For each protein, representative confocal images were shown on the left, and line scans of the relative fluorescence intensity of the protein of interest and SFPQ were shown on the right. (**G**–**J**) Bioinformatic characterization of the DNC proteome, including net charge (**G**), net charge per residue (**H**), proportion of arginine and lysine residues (**I**), and isoelectric point (pI) (**J**). The entire human proteome was used as a reference for all comparisons. The box-and-whisker plot center lines represent the median, the box edges represent the 25th and 75th percentiles, and the whiskers indicate the minimum and maximum values. Statistical significance was determined by a two-tailed unpaired Student’s *t*-test.

Using SFPQ as a bait, we obtained 79 significantly enriched protein hits (fold change ≥ 2) following AMD treatment (**Figure 3B**). Alternatively, 164 significantly enriched targets were retrieved when SRSF1 was used as the bait (**Figure 3C**). Notably, a significant proportion of protein hits (23) were identified in the proximity of both SFPQ and SRSF1 (**Figure 3D**). These proteomic hits did not overlap with previously experimentally validated^18^ DNC proteins using a targeted immunofluorescence approach, probably because of the methodological differences. The asymmetric enrichment observed between SFPQ-Turbo and SRSF1-Turbo labeling likely stemmed from the distinct subcellular distribution of SFPQ and SRSF1, as neither was exclusively DNC-localized. Integrating 13 DNC constituents that were previously experimentally validated^18^ and our proteomic results, we obtained a list of 232 proteins. Gene Ontology (GO) analysis revealed that these putative DNC proteins were closely associated with multiple aspects of RNA metabolism, including rRNA processing and RNA splicing (**Figure 3E**), implicating a potential role of DNCs in these cellular processes. We were able to confirm the enrichment of several novel proteins, such as ABT1, RCL, and SMU1, in the DNCs using immunofluorescence microscopy (**Figure 3F**)

Recent studies have demonstrated that biomolecular condensates form and sustain pH gradients at equilibrium by minimizing electrostatic repulsion among charged constituents, proposing net charge as a major determinant for the condensate pH^14,15,30^. Consistent with this idea, DNC proteins in general possessed a higher proportion of positively charged residues, including arginine and lysine, resulting in a higher total net charge (-0.6932 for the human proteome and -0.2873 for the DNC proteome) and net charge per residue (NCPR) (-0.00197 for the human proteome and -0.00084 for the DNC proteome) when compared to that of the human proteome (**Figures 3G-3I**). It has been shown that polypeptide chains buffer pH towards their pI in the dense phase^14,15,30^. We therefore analyzed the pI of the DNC-associated protein. Indeed, the median pI (7.33) of the 232 putative DNC proteins was significantly higher than that of the human proteome (median pI = 7.05) (**Figure 3J**).

Altogether, our analysis of the DNC proteome, including the identification of many previously unknown proteins, provides a foundational resource for understanding its unique pH microenvironment. We anticipate that this data will offer a valuable map for future investigations into the DNC proteomic landscape.

### SRSF1 promotes ROS synthesis through tuning the interfacial pH gradient in the DNCs

To establish the causal relationship between the interfacial pH gradient and the redox reactions, we aim to alter the DNC pH microenvironment by reducing the concentration of proteins enriched with positively charged amino acids (**Figure S5A**). Among the experimentally validated constituents of the DNCs^18^, SRSF1 possess the highest total net charge and the second highest proportion of arginine and lysine residues (**Figures S5B and S5C**). SRSF1 is a well-studied splicing factor participating in a plethora of cellular processes^31,32^. Our results have shown that the RS (arginine/serine-rich) domain of SRSF1, due to its highly positive charge, alkalizes the nucleolar microenvironment^21^. Here, we confirmed that silencing SRSF1 using a specific shRNA (**Figure S5D**) significantly obstructed the DNC alkalization when cells were stressed with AMD (**Figures 4A and 4B**). Interestingly, the reduced DNC pH was positively associated with diminished ROS concentration (**Figures 4C and 4D**). Subsequently, the interfacial potential of DNCs was also diminished when SRSF1 was depleted (**Figures 4E and 4F**).

**Figure 4.**
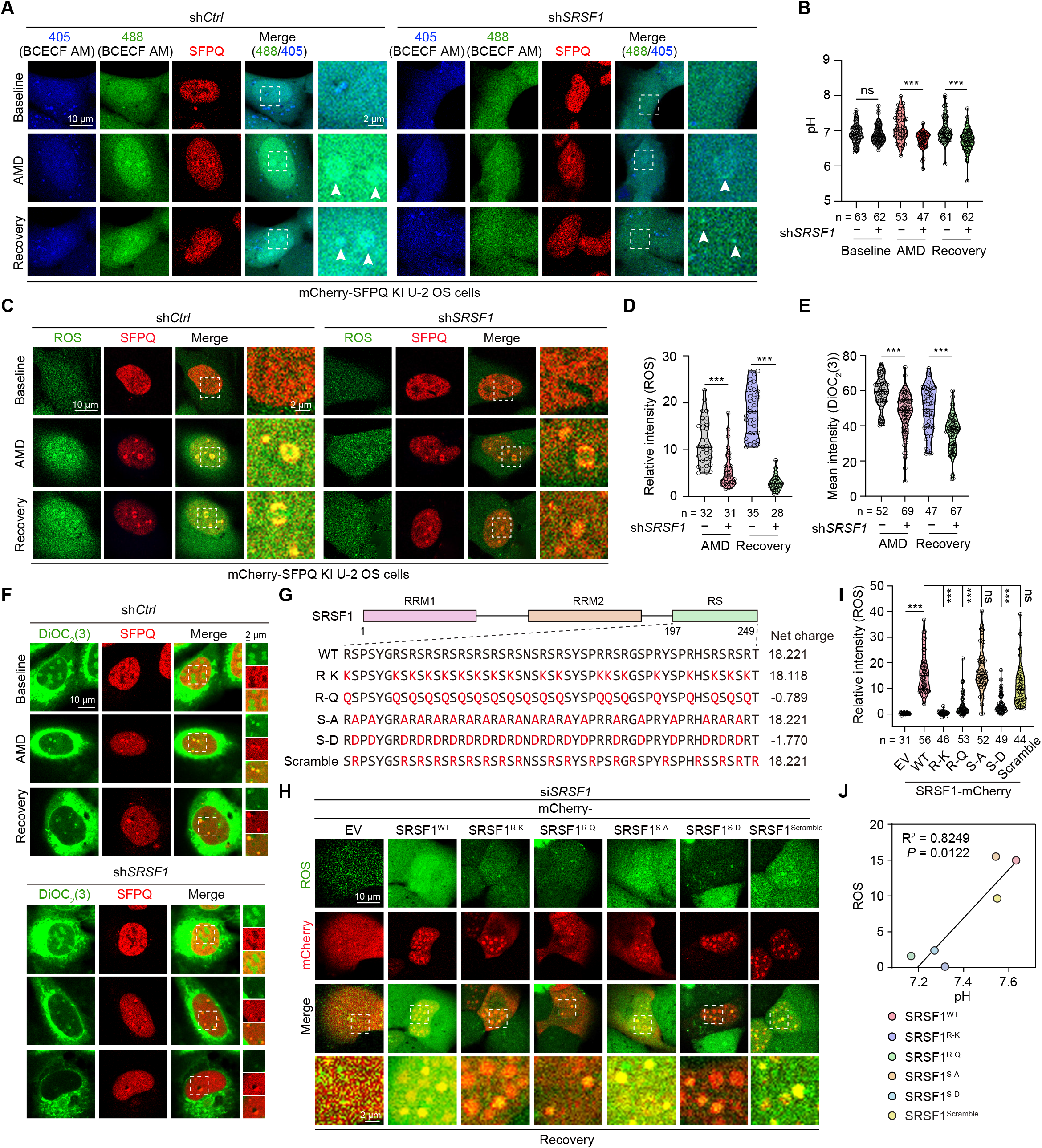
SRSF1 promotes ROS production through tuning the interfacial pH gradient in the DNCs. (**A**, **B**) Live-cell imaging of mCherry-SFPQ KI U-2 OS cells transfected with sh*Ctrl* or sh*SRSF1*, treated with AMD, and recovered under normal growth conditions in the presence of BCECF-AM. (**C**, **D**) Live-cell imaging of mCherry-SFPQ KI U-2 OS cells transfected with sh*Ctrl* or sh*SRSF1*, treated with AMD, and recovered under normal growth conditions with CM-H_2_DCFDA. (**E, F**) Live-cell imaging of mCherry-SFPQ KI U-2 OS cells transfected with sh*Ctrl* or sh*SRSF1*, treated with AMD, and recovered under normal growth conditions in the presence of DiOC_2_(3). (**G**) Schematic representation of the linear sequences SRSF1 charge mutants. (**H, I**) mCherry-SFPQ KI U-2 OS cells transfected with si*SRSF1* were reconstituted with siRNA-resistant SRSF1 variants. These cell lines were treated with AMD, and recovered under normal growth conditions. Live-cell imaging was performed using CM-H_2_DCFDA. **(J)** Linear correlation between the DNC pH and the relative ROS levels of SRSF1 WT and SRSF1 variants. For (**B**), (**D**) and (**E**), statistical significance was determined using a two-tailed unpaired Student’s *t*-test. For (**I**), statistical significance was determined using one-way ANOVA. ns, not significant; *p < 0.05; **p < 0.01; ***p < 0.001.

Despite of unclear physicochemical mechanisms, it has independently observed that the condensation of charged macromolecules triggers selective ion distribution and local pH alterations within the dense phase^9,14,30^. We designed several SRSF1 mutants to change their overall charge landscape and evaluated the subsequent impact on the DNC pH and ROS generation. Among these, all the arginine residues in the C-terminal RS domain were substituted to either lysine (referred to as SRSF1^R-K^), which retained the net positive charge but has a lower capability to engage in electrostatic interactions compared to SRSF1 WT, or to glutamine (referred to as SRSF1^R-Q^) that fundamentally abolished the net positive charge. Alternatively, we also replaced the serine residues with either alanine (referred to as SRSF1^S-A^) which presumably preserved the charge landscape, or aspartic acid (referred to as SRSF1^S-D^) to neutralize the positive charge by introducing an opposite charge (**Figure 4F**). In addition, we engineered an SRSF1 scramble mutant, in which all the arginine residues were shuffled to the neighboring location without altering the overall net charge (**Figure 4F**). Mutating these C-terminal residues did not significantly affect its protein stability (**Figure S5E**) or DNC localization (**Figures S5F and S5G**).

We then reconstituted these siRNA-resistant SRSF1 C-terminal variants into siRNA-mediated *SRSF1* knockdown cells. We carefully titrated the protein expression such that the actual protein levels of these SRSF1 mutants present in the corresponding DNCs are comparable (**Figure S5H**). As revealed by live cell imaging, WT SRSF1 robustly alkalinized the DNCs. In clear contrast, the variants with lower net charge, including SRSF1^R-Q^ (-0.789), and SRSF1^S-D^ (-1.770), failed to do so (**Figures S5I and S5J**). Note that the DNC pH was significantly lower when the SRSF1^R-K^ mutant, with a net charge slightly reduced to 18.118, compared to WT SRSF1, was reconstituted, probably due to weaker and less stable electrostatic molecular interactions initiated by lysine residues (**Figures S5I and S5J**). Strikingly, reintroducing the SRSF1^S-A^ and the scramble mutants resulted in the emergence of alkaline DNCs (**Figures S5I and S5J**). Concurrently, expression of SRSF1 mutants with lower net charge (SRSF1^R-Q^, and SRSF1^S-D^) also led to fewer redox reactions when compared to those SRSF1 variants with higher net charge (SRSF1^WT^, SRSF1^S-A^, and SRSF1^scramble^) (**Figures 4H and 4I**). We were able to establish a positive correlation between the pH values and the ROS concentrations (**Figures 4J**), implicating a strong coupling between interfacial pH gradient and redox reactions in the DNCs.

Given SRSF1’s specific role in RNA splicing, including the splicing of ribosomal protein-encoding RNAs, this function could theoretically contribute to pH modulation of the DNCs. To assess the potential contribution of abnormal RNA splicing to DNC pH modulation, we selectively silenced endogenous SRSF1 in U-2 OS cells and stably reconstituted SRSF1^WT^ and SRSF1^S-A^ mutant. RNA-sequencing analysis showed that expression of the SRSF1^S-A^ mutant moderately altered the cellular transcriptome, with 333 genes upregulated and 321 downregulated (**Figure S5K**). In stark contrast, SRSF1^S-A^ reconstitution led to widespread abnormalities in RNA alternative splicing, including more than 6700 skipped exons and other splicing errors (**Figure S5L**), which impacted core cellular processes such as cell division and the DNA damage response (**Figure S5M**). Notably, despite such profound splicing abnormalities, the SRSF1^S-A^ mutant did not appreciably alter the alkaline pH microenvironment of DNCs (**Figures S5I and S5J**). These results demonstrate that splicing dysfunction alone is not a primary mechanism driving DNC pH modulation, *per se*. Collectively, our data indicate that the high net charge of SRSF1 serves as a key determinant governing DNC pH homeostasis.

### U1-70K is required for ROS synthesis through establishing the interfacial pH gradient in the DNCs

To further consolidate the role of protein charge in dictating the pH microenvironment of the DNCs, and hence interfacial potential and ROS formation, we next focused on U1-70K, that harbors the highest proportion of positively charged residues and the second highest total net charge amongst the experimentally validated constituents of the DNCs (**Figures S5B and S5C**)^18^. U1-70K is a splicing factor involved in various key cellular pathways^33,34^. It harbors an RRM, and the rest is predicted to be disordered^35^. Although U1-70K was identified as a constituent of the DNC two decades ago^18^, its exact functions remain mysterious. First, we performed siRNA-mediated U1-70K interference (**Figure S6A**). Intriguingly, silencing U1-70K also profoundly compromised the alkalinization of the DNC (**Figures 5A and 5B**). We noticed that the lack of U1-70K caused a general decrease in the intracellular ROS levels. We therefore determined the concentration difference of ROS in the DNC and the nucleoplasm (referred to as the relative ROS concentration). As expected, loss of U1-70K also profoundly reduced the ROS production in the DNC (**Figures 5C and 5D**). Consistently, the establishment of DNC interfacial potential was also compromised in the absence of U1-70K (**Figures 5E and S6B**).

**Figure 5.**
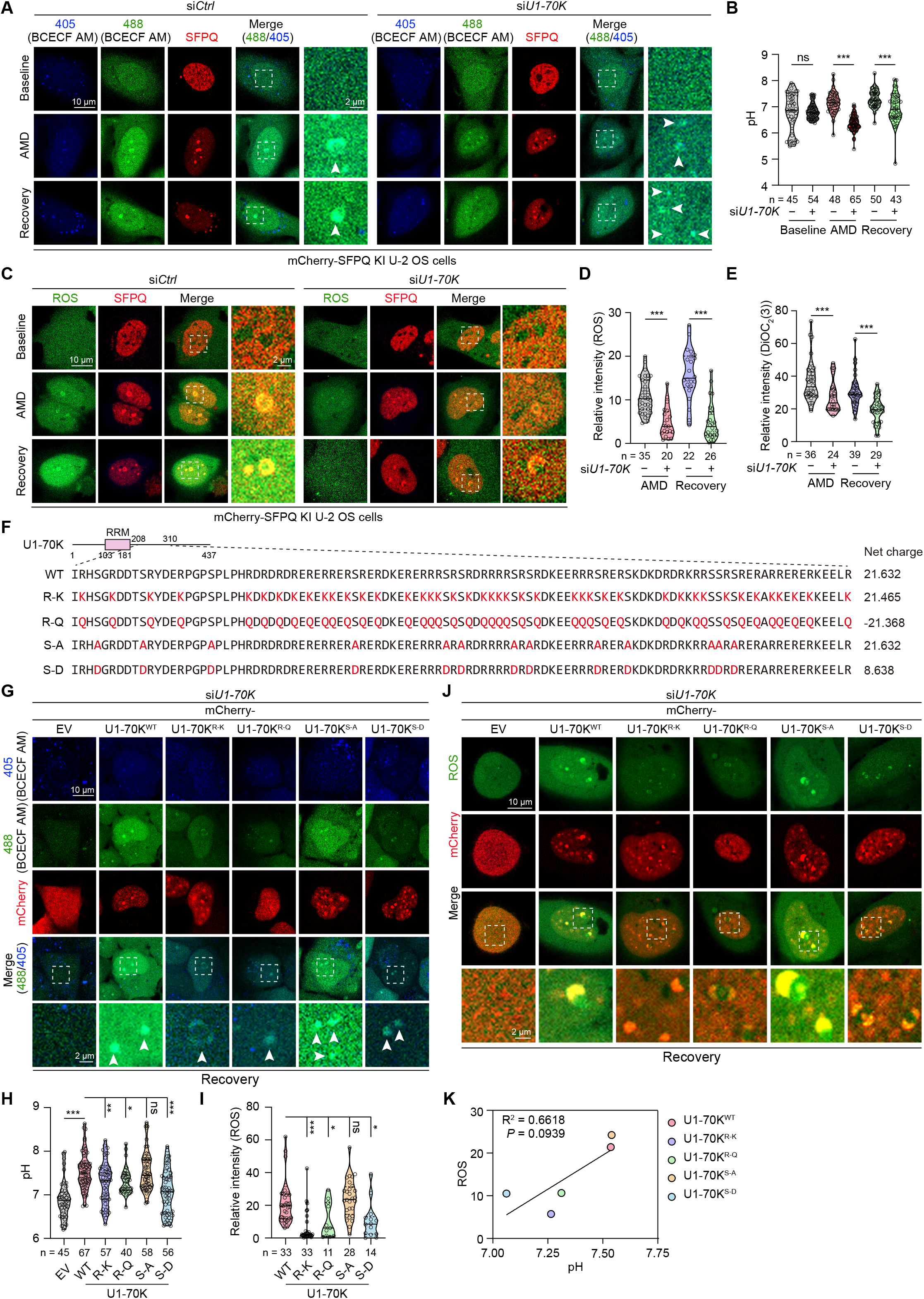
U1-70K is required for ROS synthesis through establishing the interfacial pH gradient in the DNCs. (**A-D**) mCherry-SFPQ KI U-2 OS transfected with si*Ctrl* or si*U1-70K* were treated with AMD, and recovered under normal growth conditions. Live cell imaging was performed with BCECF-AM (**A**, **B**) or CM-H_2_DCFDA (**C**, **D**). (**E**) Live-cell imaging of mCherry-SFPQ KI U-2 OS cells transfected with si*Ctrl* or sh*U1-70K*, treated with AMD, and recovered under normal growth conditions in the presence of DiOC_2_(3). (**F**) Schematic diagram of the linear sequences of U1-70K mutants. (**G-J**) WT U-2 OS stably expressing the indicated U1-70K variants were transfected with si*U1-70K*. These cells were treated with AMD, then recovered under normal growth conditions. Live cell imaging was performed with BCECF-AM (**G**, **H**) or CM-H_2_DCFDA (**I**, **J**). (**K**) Linear correlation between the DNC pH and the relative ROS levels. For (**B**), (**D**) and (**E**), statistical significance was determined using a two-tailed unpaired Student’s *t*-test. For (**H**) and (**J**), statistical significance was determined using one-way ANOVA. ns, not significant; *p < 0.05; **p < 0.01; ***p < 0.001.

When scrutinizing the charge plot of U1-70K (**Figure S6C**), we noticed a highly positively charged region (aa 208-310) that was entirely disordered (hereafter referred to as IDR), which we speculated to be involved in modulating the DNC pH. To test this, we first generated U1-70K truncation mutants with either the IDR or RRM deleted (**Figure S6D**) and examined which U1-70K region was accountable for its DNC localization. We found that deleting RRM1 was sufficient to abolish the DNC localization of U1-70K (**Figures S6E and S6F**). Therefore, this allowed us to potentially alter the charge landscape of U1-70K IDR without affecting its overall subcellular distribution in the DNCs. We then constructed several U1-70K mutants with different net charge (**Figure 5F**), including U1-70K^R-K^ (21.465), U1-70K^R-Q^ (-21.368), U1-70K^S-A^ (21.632), and U1-70K^S-D^ (8.638). We confirmed that none of the U1-70K mutants exhibited notably altered protein stability (**Figures S6G and S6H**) or aberrant DNC enrichment compared to the WT protein (**Figures S6I and S6J**). Next, we silenced the endogenous U1-70K using siRNA and transiently expressed these U1-70K mutants that were resistant to siRNA interference, and performed live cell imaging to quantify the pH and ROS formation in the DNCs. We again titrated the protein expression such that the actual protein levels of these U1-70K mutants present in the corresponding DNCs are comparable (**Figure S6K**). Indeed, arginine substitution, either to lysine or glutamine, significantly reduced U1-70K’s potential to alkalize the DNCs, so did arginine to aspartic acid substitution (**Figures 5G and 5H**). Strikingly, expression of the U1-70K^S-A^ mutant resulted in alkaline DNCs and ROS formation (**Figures 5G-5J**). Again, we established a positive correlation between the pH values and the relative ROS concentrations (**Figure 5K**), strengthening the idea that interfacial pH gradient and redox reactions are coupled in the DNCs, which is regulated by the characteristic net charge of proteins, such as SRSF1 and U1-70K.

### DNC-derived ROS modulates the biophysical properties of redox-reactive DNC constituents

Given that ROS is highly reactive with biological macromolecules, we next sought to delineate the impact of accumulated ROS on the biophysical properties of the DNC proteins. Since ROS, H_2_O_2_ in particular, are prone to reversibly oxidize critical, redox-sensitive cysteine and methionine residues on the target proteins, we analyzed the proportion of cysteine and methionine residues of the experimentally validated DNC proteins. We selected DDX5 and SFPQ for the subsequent investigation due to their relatively overrepresented cysteine and methionine residues, respectively (**Figures 6A and 6B**), and the fact that SFPQ redox state mediates stress granule dissolution^36^. Meanwhile, we chose SRSF1 and U1-70K for comparison because of their relatively lower abundance of redox-sensitive residues. Next, we tagged the endogenous loci of *DDX5* and *U1-70K* with the mCherry coding sequence in U-2 OS cells (**Figures S7A and S7B**). Considering that protein oxidation often alters the biophysical properties of target proteins^36,37^, we first performed fluorescence recovery after photobleaching (FRAP) to assess the effect of ROS on the molecular exchange of DDX5 and SFPQ. Remarkably, removing the excessive ROS in the DNCs with NAC significantly accelerated the fluorescence recovery rate of both DDX5 and SFPQ at the endogenous levels (**Figures 6C-6F**). This was true even for the exogenous GFP-tagged DDX5 (**Figures S7C and S7D**). In striking contrast, the presence of ROS had no appreciable impact on the molecular diffusivity of either SRSF1 or U1-70K (**Figures S7E-S7H**). This finding prompted us to examine whether the selective partition of proteins within the DNC dense phase is regulated by oxidation. Notably, NAC treatment reduced the enrichment of DDX5 and SFPQ within the DNCs, leading to a lower partition coefficient of both proteins (**Figures 6G-6J**). Consistently, the partition of either SRSF1 or U1-70K in the DNC was not appreciably altered by the presence or absence of ROS (**Figures S7I-S7L**).

**Figure 6.**
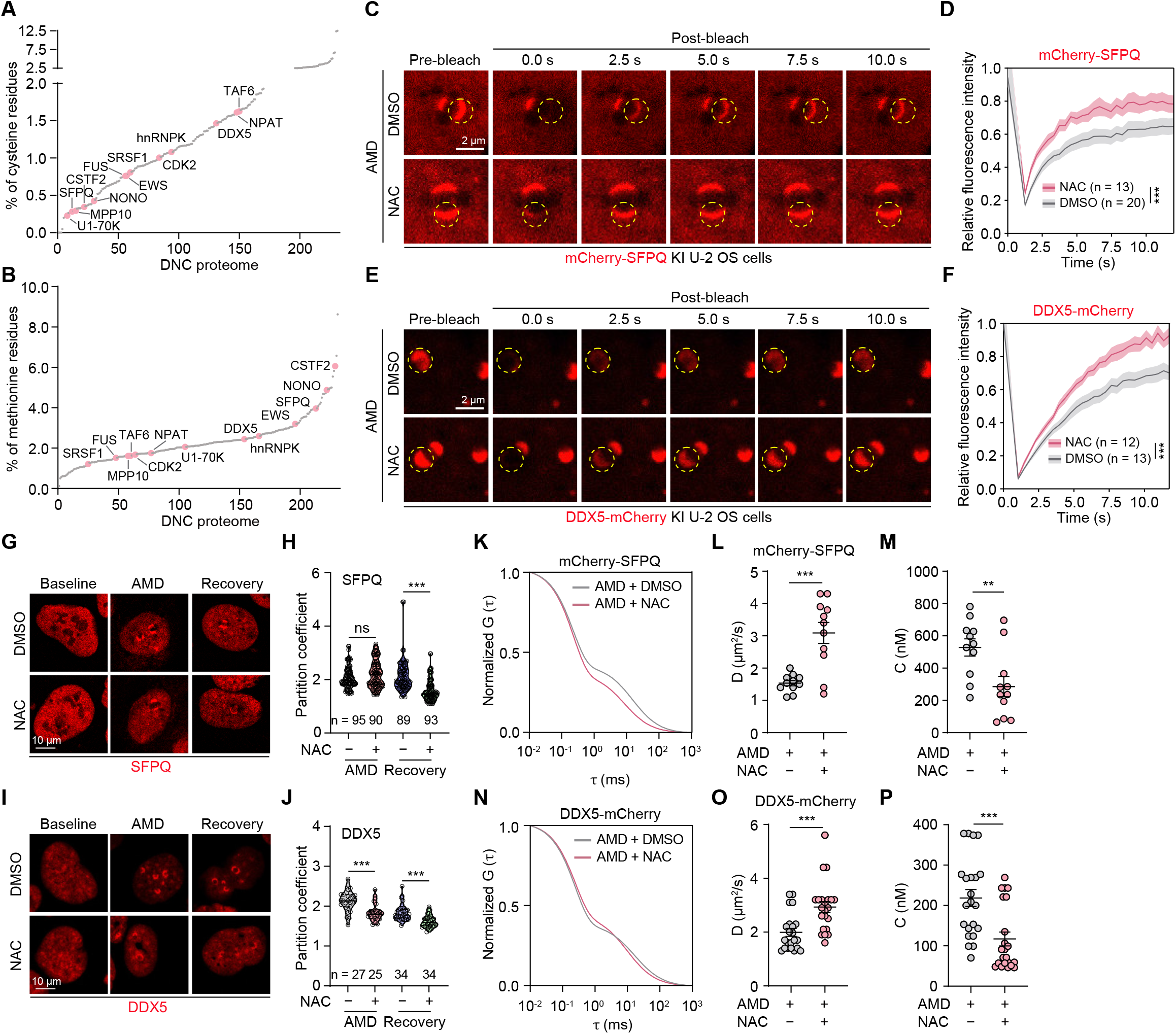
DNC-derived ROS modulates the biophysical properties of redox-reactive DNC constituents. (**A**, **B**) Bioinformatics analyses of the proportion of cysteine (**A**) and methionine (**B**) residues of the putative DNC proteome. (**C**-**F**) U-2 OS cells from DMSO and NAC-treated groups were subjected to a 4-hour AMD treatment, followed by a 2-hour recovery. FRAP was then performed to monitor and quantitatively analyze the time-dependent changes in fluorescence intensity of mCherry-SFPQ (**C, D**) or DDX5-mCherry (**E, F**). Data are presented as mean ± SD. ns, not significant; ***p < 0.001 by two-way ANOVA). (**G**-**J**) U-2 OS cells pretreated with DMSO and NAC were exposed to AMD for 4 hours followed by 2-hour recovery. The partition coefficient was quantified as the ratio of the mean fluorescence intensity in the DNC to that in the nucleoplasm for SFPQ (**G**, **H**) and DDX5 (**I**, **J**). ns, not significant; ***p < 0.001 by a two-tailed unpaired Student’s *t*-test. **(K**-**N)** Normalized autocorrelation functions G(τ) obtained from FCS measurements of mCherry–SFPQ (**K**) and DDX5–mCherry (**N**) in U-2 OS cells treated with AMD + DMSO or AMD + NAC. Scatter plots showing the diffusion coefficients (D) derived from FCS fitting for mCherry–SFPQ (**L**) and DDX5–mCherry (**O**). Treatment with AMD + NAC significantly increased D compared with AMD + DMSO (D = 1.5 ± 0.3 vs. 3.1 ± 1.1 μm^2^/s for mCherry–SFPQ; D = 2.0 ± 0.7 vs. 3.1 ± 1.1 μm^2^/s for DDX5–mCherry; n = 10 cells per condition). Apparent cellular concentrations (C) of mCherry–SFPQ (**M**) and DDX5-mCherry (**P**) were determined from FCS measurements. NAC treatment markedly reduced C compared with AMD + DMSO (C = 528 ± 176 vs. 286 ± 208 nM for mCherry– SFPQ; C = 228 ± 95 vs. 99 ± 69 nM for DDX5-mCherry; n = 10 cells per group). Data are shown as mean ± SD. **p < 0.01; ***p < 0.001 by a two-tailed unpaired Student’s *t*-test.

Next, we used fluorescence correlation spectroscopy (FCS) technique to quantitatively determine the molecular concentration and diffusion coefficient of the DDX5 and SFPQ in the DNCs at single molecule level. The diffusion coefficient for mCherry-SFPQ in the DNCs was around 1.53 micrometer^2^/s, which was increased to 3.09 micrometer^2^/s when the ROS was depleted (**Figures 6K and 6L**). In addition, ROS elimination reduced the molecular concentration of mCherry-SFPQ in the DNCs from 527.7 nM to 285.7 nM (**Figure 6M**), strengthening the aforedescribed findings that the selective partition of redox-responsive proteins within the DNCs is regulated by ROS. Similar conclusions were reached when the FCS experiments were performed with DDX5-mCherry KI U-2 OS cells (**Figures 6N-6P**). We then performed sucrose gradient centrifugation to examine if the oligomeric status of the DNC proteins is affected by ROS (**Figure S8A**). In fact, the overall migration pattern of the DNC proteins was not appreciably influenced by ROS (**Figure S8B**), suggesting that changes to the biophysical properties of the DNC proteins by ROS likely occur only to restricted locations (i.e. DNCs).

### Eliminating ROS accelerates the structural and functional restoration of the nucleolus

To elucidate the role of ROS in the nucleolar organization and functionality, we took advantage of two widely used ROS scavengers, NAC and GSH (**Figure 7A**). Cells were treated with a lower dose of AMD (0.1 µM) to disrupt the nucleolar organization and allowed for functional recovery for an extended time, in the presence of NAC. We found that removing ROS notably expedited the size growth of GC (NPM1^+^), and DFC (FBL^+^), but not that of UBF^+^ FC during cellular recovery from AMD treatment (**Figures 7B-7E, S9A, and S9B**). Given that the primary function of the nucleolus is to promote ribosomal biogenesis, we therefore performed a fluorescence in situ hybridization (FISH) experiment using a probe for 5’-ETS, and an immunostaining procedure using an antibody specific against mature rRNA to assess rRNA processing and maturation, respectively. We observed that scavenging the DNC ROS prominently accelerated both rRNA processing and maturation during the functional recovery of the nucleolus from AMD stress (**Figures 7F-7I**). Because AMD exposure elicits DNA damage and the nucleolus is closely related to DNA repair, we immunostained the cells with antibodies selectively against γH2AX to visualize the foci of DNA damage as a potential indirect assessment of the nucleolar function. We found that the DNA repair was much more efficient when cellular ROS was removed (**Figures 7J and 7K**). This was also in line with the observation that cellular p53 levels that were readily induced by DNA damage, were markedly lower in cells upon ROS removal (**Figures 7L and 7M**). Similarly, eradicating cellular ROS by GSH also evidently accelerated rRNA processing and maturation, and DNA repair (**Figures S9C-S9H**). Collectively, our data imply that removing the excessive ROS that is produced by the interphase electric field in the DNC improved cellular resilience and homeostasis.

**Figure 7.**
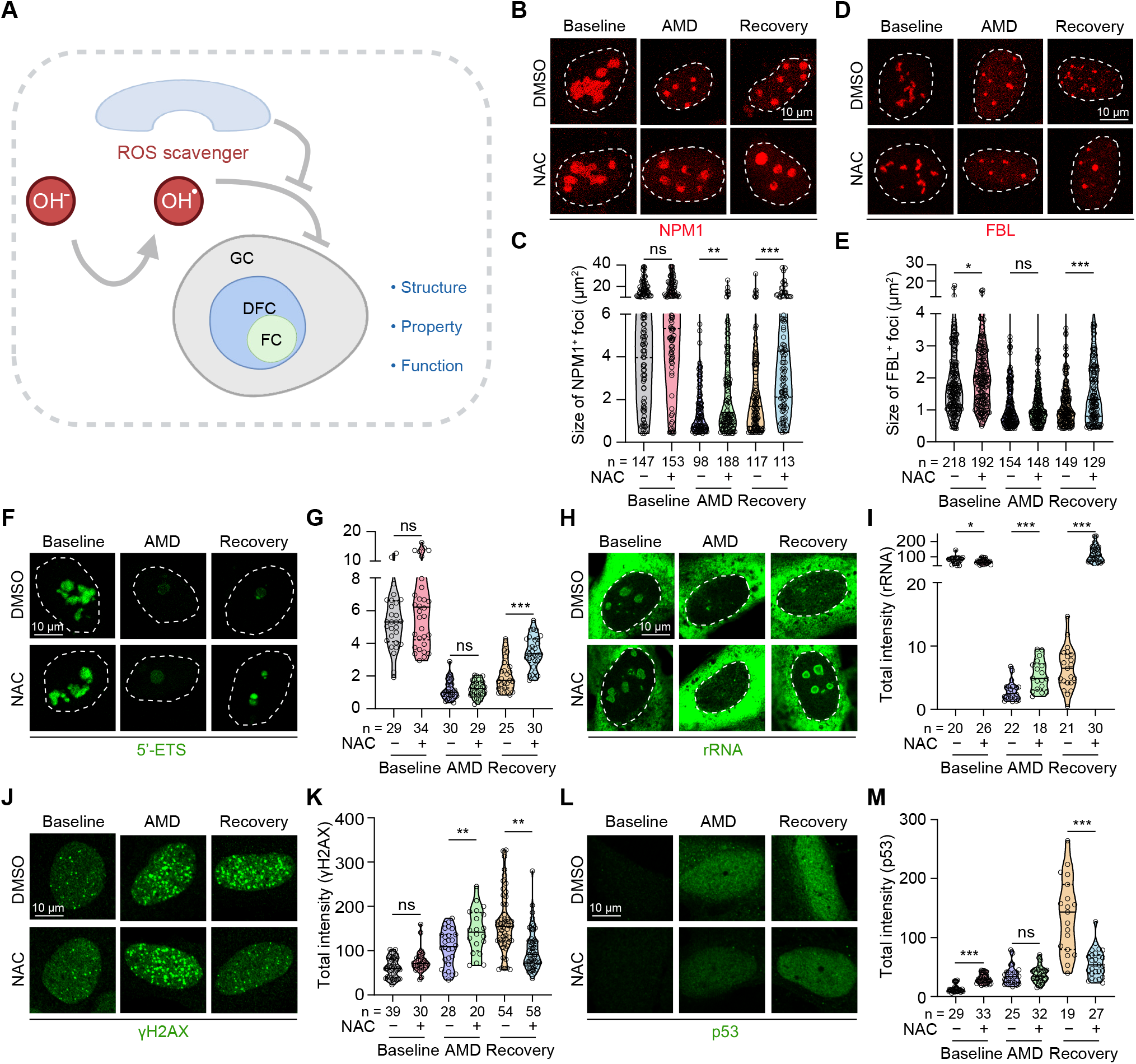
Eliminating ROS accelerates the structural and functional restoration of the nucleolus. (**A**) Schematic illustration of the experimental design. (**B-E**) After treating U-2 OS cells (pretreated with DMSO or NAC) with AMD for 4 hours, these cells were recovered for 24 hours. Cells were subjected to immunostaining using the indicated antibodies. The size of NPM1 (**B**, **C**) or FBL (**D**, **E**) foci was quantified. (**F**-**M**) After treating U-2 OS cells (pretreated with DMSO or NAC) with AMD for 4 hours, these cells were recovered for 24 hours. The cells were processed for either FISH using a probe for 5’-ETS (**F**, **G**), or immunostaining using antibodies against rRNA (**H**, **I**), γH2AX (**J**, **K**), and p53 (**L**, **M**), respectively. The total intensity of 5’-ETS (**G**), rRNA (**I**), γH2AX (**K**), and p53 (**M**) were quantified, respectively. For (**C**), (**E**), (**G**), (**I**), (**K**) and (**M**), statistical significance was determined using a two-tailed unpaired Student’s *t*-test. ns, not significant; *p < 0.05; **p < 0.01; ***p < 0.001.

## Discussion

Our study, utilizing nucleolar caps as a model system for the interfacial microenvironment of phase-separated biomolecular condensates, reveals a non-canonical, enzyme-independent pathway for ROS generation in mammalian cells, driven fundamentally by electrochemical potential gradients. This finding challenges the prevailing paradigm that predominantly attributes intracellular ROS production to dedicated enzymatic machinery, such as the mitochondrial electron transport chain or NADPH oxidases. Instead, we propose a model where the physicochemical properties inherent to the interfaces of biomolecular condensates can directly potentiate redox chemistry. The coupling of interfacial electric fields, ion selectivity, and distinct physicochemical milieus between co-existing phases creates a unique platform for non-enzymatic chemical activity, a mechanism we posit is not exclusive to nucleolar caps but likely operational at the boundaries of other condensates, such as stress granules, P bodies, and nuclear speckles, across diverse physiological and pathological contexts (**Figure S9I**).

The mechanistic foundation of this phenomenon appears to lie in the ability of intense interfacial electric fields to significantly alter the effective ionization energy of solvated molecules and modulate local dielectric constants. This restructuring of the energy landscape can lower activation barriers for redox reactions and steer reaction pathways in a manner distinct from bulk solution or homogeneous cellular compartments. The electrochemical potential gradient established by the asymmetric partitioning of ions and charged biomolecules between the dense and dilute phases provides a driving force for spontaneous electron and proton transfer. Consequently, the generation of ROS we observed can be rationalized by the facilitated formation and stabilization of radical intermediates at these dynamic interfaces, where factors such as localized pH shifts, ionic strength, and water activity differ markedly from the surrounding nucleoplasm.

This discovery compels a re-evaluation of the functional repertoire of biomolecular condensates, positioning them not merely as passive organizational hubs but as active contributors to cellular metabolism and signaling through non-enzymatic chemical catalysis. The nucleolar cap, formed during transcriptional stress, may thus serve a previously unrecognized role in modulating the cellular redox state. The enzyme-free generation of ROS at such interfaces could represent a rapid, spatially constrained response to metabolic demands or stress signals, potentially influencing processes like rRNA processing arrest, DNA damage response, and cell fate decisions. This paradigm of “metabolons” for non-enzymatic chemistry expands our understanding of how cells leverage phase separation to compartmentalize and control fundamental biochemical processes.

Importantly, the dysregulation of this condensate-mediated ROS generation presents a plausible pathophysiological mechanism, particularly in neurodegenerative diseases. Many proteins that form pathogenic aggregates in conditions such as amyotrophic lateral sclerosis and frontotemporal dementia (e.g., FUS, TDP-43) are known to undergo aberrant phase separation^38,39^. Our model suggests that the persistent, misfunctioning interfaces of these pathological condensates could become chronic, unregulated sources of ROS. This sustained oxidative stress would exacerbate proteotoxicity, promote further protein misfolding, and ultimately contribute to the vicious cycle of neuronal damage and loss. Thus, the active chemical nature of these condensates, when dysregulated, transforms an organizational platform into a driver of cellular vulnerability.

In conclusion, our work elucidates a fundamental mechanism by which biomolecular condensates directly engage in redox chemistry through the physicochemical properties of their interfaces. This enzyme-free, electrochemically driven ROS generation underscores the multifaceted roles of phase separation in cellular physiology and pathology. Harnessing or mitigating this chemical activity at condensate interfaces may open new therapeutic avenues for diseases characterized by oxidative stress and aberrant phase separation, from neurodegeneration to cancer.

### Limitations of the study

While our findings provide compelling evidence for a novel mechanism driving redox reactions, several open questions remain for future investigation. First, systematically manipulating condensate composition-varying salt types, pH, and the presence of specific redox-active molecules-will be essential to delineate the precise chemical rules governing this phenomenon and to establish causal relationships among pH, interfacial potential, and redox reactions. Second, we have not fully excluded the potential contribution of aberrant splicing to ROS generation. Third, the impact of ROS on biomolecular constituents of condensates is largely based on cellular correlation; direct biochemical evidence from *in vitro* reconstitution experiments would be ideal. Finally, more physiologically relevant models could illuminate the spatiotemporal dynamics and physiological scope of this process in live cells.

## Resource availability

### Lead contact

Further information and requests for resources and reagents should be directed to and will be fulfilled by the lead contact, Bo Wang.

### Materials availability

Materials used in this study will be made available upon reasonable request to the lead contact.

### Data and code availability

- The RNA-sequencing data used in this study have been deposited to Gene Expression Omnibus and are publicly available as of the date of publication (GEO: GSE328521). Proteomics data have been deposited in ProteomeXchange and are publicly available as of the date of publication (PXD070934). Raw imaging data of microscopy, gels, and blots have been deposited at Mendeley Data and are publicly available as of the date of publication (DOI: 10.17632/vrs925kdny.1).
- This paper does not report original code
- Any additional information required to reanalyze the data reported in this work paper is available from the lead contact upon request

## Acknowledgements

All the imaging data were acquired in the Core Facility of Biomedical Sciences, Xiamen University. This work was supported by the National Natural Science Foundation of China (32470726, and 32522024 to B.W.; 22304184 to J.G; 22494700, and 22477102 to X.Z.; 325B2024 to J.C.), the Natural Science Foundation of Fujian Province, China (2024J010004 to B.W.), Shenzhen Science and Technology Program (JCYJ20250604122935047 to B.W.), the Fundamental Research Funds for the Central Universities (B.W.), the Guangdong Talents Program (2023QN10Y004 to J.G.) and CAS Pioneer Hundred Talents Program (J.G.).

## Author contributions

B.W. and S.M. designed the experiments. S.M., X.Z., Z.M., F.W., J.C., J.G., W.L., C.Z., Y.Z., X.Z., J.G., and B.W. performed experiments and/or analyzed the data. B.W. wrote the initial draft. J.G. edited the manuscript.

## STAR★Methods

### Key resources table

| REAGENT or RESOURCE | SOURCE | IDENTIFIER |
|---|---|---|
| Antibodies |  |  |
| mouse anti-Flag | GNI | Cat: #GNI4110-FG; RRID: AB_3067995 |
| rabbit anti-U1-70K | Abcam | Cat: #ab83306; RRID: AB_10673827 |
| mouse anti-SRSF1 | Invitrogen | Cat: #32-4500; RRID: AB_2533079 |
| rabbit anti-SFPQ | Proteintech | Cat: #15585-1-AP; RRID: AB_10697653 |
| mouse anti-p53 | Proteintech | Cat: #60283-1-Ig; RRID: N/A |
| mouse anti-FUS | Proteintech | Cat: #68262-1-Ig; RRID: AB_2935347 |
| mouse anti-FBL | Proteintech | Cat: #66985-1-Ig; RRID: AB_2882303 |
| rabbit anti-NCL | Proteintech | Cat: #10556-1-AP; RRID: AB_2082423 |
| rabbit anti-FBL | Proteintech | Cat: #16021-1-AP; RRID: AB_2105788 |
| mouse anti-UBF | Santa Cruz Biotechnology | Cat: #SC-13125; RRID: AB_671403 |
| mouse anti-NPM1 | Invitrogen | Cat: #32-5200; RRID: AB_2533084 |
| rabbit anti-α-Tubulin | Cell Signaling Technology | Cat: #2125S; RRID: AB_2619646 |
| mouse anti-rRNA | Santa Cruz Biotechnology | Cat: #SC-33678; RRID: AB_628226 |
| rabbit anti-γH2AX | Abclonal | Cat: #AP0687; RRID: AB_2863808 |
| Horseradish peroxidase-conjugated Goat Anti-Rabbit IgG (H+L) | JacksonImmunoResearch | Cat: #111-035-003; RRID: AB_2313567 |
| Horseradish peroxidase-conjugated Goat Anti-Mouse IgG (H+L) | JacksonImmunoResearch | Cat: #115-035-003; RRID: AB_10015289 |
| AlexaFluor 488AffiniPureGoatAnti-MouseIgG | JacksonImmunoResearch | Cat: #115-545-003; RRID: AB_2338840 |
| AlexaFluor 594AffiniPureGoatAnti-MouseIgG | JacksonImmunoResearch | Cat: #115-585-003; RRID: AB_2338871 |
| AlexaFluor 647AffiniPureGoatAnti-MouseIgG | JacksonImmunoResearch | Cat: #115-605-003; RRID: AB_2338902 |
| Streptavindin-HRP | APExBIO | Cat: #K1229; RRID: N/A |
| rabbit anti-Vinculin | Proteintech | Cat: #26520-1-AP; RRID: AB_2868558 |
| rabbit anti-DDX5 | Diagbio | Cat: #db11221; RRID: N/A |
| mouse anti-NPAT | Abcam | Cat: #ab70595; RRID: AB_1269585 |
| rabbit anti-mCherry | Proteintech | Cat: #26765-1-AP; RRID: AB_2876881 |
| Bacterial and virus strains |  |  |
| DH5α Escherichia coli strain | AlpalifeBio | Cat. #KTSM101L |
| Chemicals, peptides, and recombinant proteins |  |  |
| TritonX-100 | Biotopped | Cat: #T6200G |
| Tween-20 | BioFroxx | Cat: #1247LT001 |
| DNA marker (2000bp) | LABLEAD | Cat: #L2000 |
| DNA marker (1kb) | LABLEAD | Cat: #L1000P |
| Diethyl pyrocarbonate | MCE | Cat: #HY-D0846 |
| DAPI | Thermo Fisher | Cat: #D1306 |
| Doxorubicin hydrochloride (DRB) | MCE | Cat: #HY-15142 |
| Carbonyl cyanide 3-chlorophenylhydrazone (CCCP) | MCE | Cat: #HY-100941 |
| N-Acetyl-L-cysteine (NAC) | MCE | Cat: #HY-B0215 |
| L-Glutathione reduced (GSH) | MCE | Cat: #HY-D0187 |
| L-carnosine | Targetmol | Cat: #T4732 |
| Actinomycin D (AMD) | MCE | Cat: #HY-17559 |
| 30% SDS | Leagene | Cat: #R00175 |
| 30% acrylamide | LABLEAD | Cat: #A3291 |
| Agarose | LABLEAD | Cat: #AG0100 |
| Super GelBlue<sup>TM</sup> | UElandy | Cat: #S2019L |
| Pentafluorobenzenesulfonyl fluorescein | Amgicam | Cat: #ajci73430 |
| DiOC<sub>2</sub>(3) | Maokangbio | Cat: #MX4008-100MG |
| CM-H<sub>2</sub>DCFDA | Invitrogen | Cat: #C6827 |
| BCECF-AM | Beyotime | Cat: #S1006 |
| RNase A | LABLEAD | Cat: #R4875 |
| RNase I | Thermo Fisher | Cat: #EN0601 |
| RNase T1 | Beyotime | Cat: #R7096S |
| RNase III | Beyotime | Cat: #R7086S |
| Puromycin | LABLEAD | Cat: #P067 |
| 4% paraformaldehyde | Leagene | Cat: #DF0135 |
| Ethanol absolute | Sinopharm | Cat: #10009218 |
| Isopropanol | Sinopharm | Cat: #80109218 |
| Yeast tRNA | ACMEC biochemical | Cat: #T48660 |
| Bovine Serum Albumin | LABLEAD | Cat: #0332 |
| Glycine | LABLEAD | Cat: #GAS10 |
| Nigericin | Shanghai Yuanye Bio-Technology | Cat: #S25116 |
| NaCl | LABLEAD | Cat: #X190 |
| Sucrose | LABLEAD | Cat: #S0334-1 |
| Critical commercial assays |  |  |
| HiScript IV All-in-One Ultra RT SuperMix | Vazyme | Cat: #R433-01 |
| AG RNAex Pro Reagent | Accurate Biotechnology | Cat: #AG21102 |
| Qubit™ RNA Broad-Range Assay Kit | Thermo Fisher | Cat: #Q10210 |
| 2× Universal SYBR Green Fast qPCR mix | Abclonal | Cat: #RM21203 |
| 2× Phanta Max Master Mix | Vazyme | Cat: #P515-01 |
| Polybrene | Santa Cruz Technology | Cat: #sc-134220 |
| Fluoromount-G Mounting medium | SouthernBiotech | Cat: #0100-01 |
| Immobilon Western HRP Substrate | Merck Millipore | Cat: #WBKLS0050 |
| DMEM medium | BasalMedia | Cat: #L120KJ |
| Gibco™ DMEM/F-12, no phenol red | Gibco | Cat: #21-041-025 |
| McCoy's 5A medium | BasalMedia | Cat: #L630KJ |
| Fetal bovine serum | Moybio | Cat: #S450 |
| Penicillin/streptomycin 100X | BasalMedia | Cat: #S110JV |
| 200 mM L-glutamine solution | BasalMedia | Cat: #S210JV |
| Lipofectamine RNAiMax | Thermo Fisher | Cat: #13778150 |
| Streptavidin-conjugated magnetic beads | GenScript | Cat: #L00936 |
| Neofect™ DNA transfection reagent | Neofect (beijing) biotech | Cat: #TF201201 |
| 2× MultiF Seamless Assembly Kit | Abclonal | Cat: #RK21020 |
| Deposited data |  |  |
| Mass spectrometry data | This paper | ProteomeXchange: PXD070934: https://www.iprox.cn//page/SCV017.html?query=PXD070934 |
| Raw RNA sequencing data | This paper | Gene Expression Omnibus: GSE328521 |
| Raw imaging data of microscopy, gels, and blots | This paper | Mendeley Data: https://doi.org/10.17632/vrs925kdny.1 |
| Experimental models: Cell lines |  |  |
| Human: U-2 OS | ATCC | ATCC HTB-96 |
| Human: HEK 293T | ATCC | ATCC CRL-1573 |
| Human: HeLa | ATCC | ATCC CCL-2 |
| Human: Huh7 | ATCC | ATCC PTA-8561 |
| Human: HCT 116 | ATCC | ATCC CCL-247 |
| Oligonucleotides |  |  |
| shRNA, siRNA, sgRNA, and RT-PCR primers | This paper | See Table S1 |
| Recombinant DNA |  |  |
| pcDNA3.3-Flag-Halo-TCOF1 | This paper | N/A |
| pcDNA3.3-Flag-Halo-SRSF1 | This paper | N/A |
| pcDNA3.3-Flag-Halo-SFPQ | This paper | N/A |
| pcDNA3.3-Flag-RRM<sub>SRSF1</sub>-roGFP1 | This paper | N/A |
| pcDNA3.3-Flag-BFP-RRM<sub>SRSF1</sub> | This paper | N/A |
| pcDNA3.3-Flag-BFP-RRM<sub>SRSF1</sub>-MT1A | This paper | N/A |
| pcDNA3.3-Flag-BFP-RRM<sub>SRSF1</sub>-Catalase | This paper | N/A |
| pcDNA3.3-Flag-RCL | This paper | N/A |
| pcDNA3.3-Flag-ABT1 | This paper | N/A |
| pcDNA3.3-Flag-SMU1 | This paper | N/A |
| pcDNA3.3-Flag-mCherry-SRSF1<sup>WT</sup> | This paper | N/A |
| pcDNA3.3-Flag-mCherry-SRSF1<sup>R-K</sup> | This paper | N/A |
| pcDNA3.3-Flag-mCherry-SRSF1<sup>R-Q</sup> | This paper | N/A |
| pcDNA3.3-Flag-mCherry-SRSF1<sup>S-A</sup> | This paper | N/A |
| pcDNA3.3-Flag-mCherry-SRSF1<sup>S-D</sup> | This paper | N/A |
| pcDNA3.3-Flag-mCherry-SRSF1<sup>Scramble</sup> | This paper | N/A |
| pcDNA3.3-Flag-mCherry-U1-70K<sup>WT</sup> | This paper | N/A |
| pcDNA3.3-Flag-mCherry-U1-70K<sup>R-K</sup> | This paper | N/A |
| pcDNA3.3-Flag-mCherry-U1-70K<sup>R-Q</sup> | This paper | N/A |
| pcDNA3.3-Flag-mCherry-U1-70K<sup>S-A</sup> | This paper | N/A |
| pcDNA3.3-Flag-mCherry-U1-70K<sup>S-D</sup> | This paper | N/A |
| pcDNA3.3-Flag-mCherry-U1-70K<sup>ΔIDR</sup> | This paper | N/A |
| pcDNA3.3-Flag-mCherry-U1-70K<sup>ΔRRM</sup> | This paper | N/A |
| pUC19-mCherry-SFPQ | This paper | N/A |
| pUC19-mCherry-NPM1 | This paper | N/A |
| pUC19-FBL-mCherry | This paper | N/A |
| pUC19-mCherry-TCOF1 | This paper | N/A |
| pUC19-DDX5-mCherry | This paper | N/A |
| pUC19-GFP-SRSF1 | This paper | N/A |
| pUC19-U1-70K-mCherry | This paper | N/A |
| pCW57.1-Flag-SFPQ-TurBoID | This paper | N/A |
| pCW57.1-Flag-SRSF1<sup>WT</sup> | This paper | N/A |
| pCW57.1-Flag-SRSF1<sup>R-K</sup> | This paper | N/A |
| pCW57.1-Flag-SRSF1<sup>R-Q</sup> | This paper | N/A |
| pCW57.1-Flag-SRSF1<sup>S-A</sup> | This paper | N/A |
| pCW57.1-Flag-SRSF1<sup>S-D</sup> | This paper | N/A |
| pLV-Flag-GFP-DDX5 | This paper | N/A |
| pLV-Flag-SRSF1-TurBoID | This paper | N/A |
| pLKO.1 | Addgene | Cat# 10878; RRID: Addgene_10878 |
| px459 | Addgene | Cat# 101716; RRID: Addgene_101716 |
| psPAX2 | Addgene | Cat# 12260; RRID: Addgene_12260 |
| pMD2.G | Addgene | Cat# 12259; RRID: Addgene_12259 |
| pUC19 | Addgene | Cat# 50005; RRID: Addgene_50005 |
| Software and algorithms |  |  |
| SnapGene | SnapGene | https://www.snapgene.com/ |
| GraphPad Prism 10 | GraphPad Software | https://www.graphpad.com |
| Fiji (ImageJ) | ImageJ | https://fiji.sc/ |
| Python code for charge analysis | Yamazaki et al., 2022<sup>40</sup> | https://www.python.org/ |
| PONDR | N/A | http://www.pondr.com/ |

## Experimental model and study participant details

### Cell line

The U-2 OS (HTB-96), HeLa (CCL-2), HEK293T (CRL-1573), HuH7 (PTA-8561) and HCT 116 (CCL-247) cell lines were derived from the American Type Culture Collection (ATCC). Cells were cultured in DMEM medium (BasalMedia; L120KJ), except for the U-2 OS cell line, which was maintained in McCoy’s 5A medium (BasalMedia; L630KJ). Cells were cultured in medium containing 10% fetal bovine serum (Moybio; S450), penicillin/streptomycin (BasalMedia; S110JV) and Glutamax (BasalMedia; S210JV) at 37°C in a 5% CO_2_ cell culture incubator. All cell lines were free of mycoplasma contamination.

## Method details

### Plasmid Construction

Vector backbones were linearized via digestion with restriction enzymes. Target fragments were amplified using the high-fidelity DNA polymerase 2×Phanta Max Master Mix (Vazyme; P515-01). PCR primers were designed using SnapGene software, with 15 bp homologous arms. The targeted plasmids were constructed via seamless cloning using the 2×MultiF Seamless Assembly Kit (Abclonal; RK21020), and Sanger sequencing was subsequently performed to verify the correctness of the plasmid sequence.

For constructing the plasmids used in rescue assays, SRSF1 and U1-70K mutants were inserted into the pC3.3 vector. To generate a plasmid for silencing SRSF1 expression, a short hairpin RNA (shRNA) targeting *SRSF1* (5’-TGGTCGCGACGGCTATGATTA-3’) was cloned into the pLKO.1 vector.

### Transfection, lentivirus infection and drug treatment

For transient protein expression, Neofect™ DNA transfection reagent (Neofect (beijing) biotech; TF201201) was utilized. For the construction of stable expression cell lines, lentiviruses were acquired from co-transfection of the plasmid system by Lipofectamine 2000 in HEK293T cell lines, including cloned plasmids, psPAX2 (Addgene; 12260), and pMD2.G (Addgene; 12259). 48 hours after the transfection, the supernatant contain the viral particles was collected and filtered, and cells were infected for 24 hours in the presence of 8 µg/mL polybrene (Santa Cruz Technology; sc-134220).

Chemical compounds, dissolved in DMSO or PBS, were directly added to cells as follows: 1 μM Actinomycin D (AMD) (MCE; HY-17559) for 4 hours; 2 μM doxorubicin (DRB) (MCE; HY-15142) for 6 hours; 7.5 μM Carbonyl cyanide 3-chlorophenylhydrazone (CCCP) (MCE; HY-100941) for 30 min; L-carnosine (Targetmol; T4732) for 4 hours; 1 mM N-Acetyl-L-cysteine (NAC) (MCE; HY-B0215) for 6 hours; 0.5 M L-Glutathione reduced (GSH) (MCE; HY-D0187) for 6 hours.

### RNA interference

Cells were transfected with the indicated siRNAs using Lipofectamine RNAiMAX (Thermo Fisher; 13778150) according to the manufacturer’s instructions and analyzed at least 48 hours after transfection. The siRNAs synthesized by GenePharma (Shanghai, China) were used as follows: si*U1-70K*, 5’-UGAUAGGUCCGUACACCUCTT-3’ and 5’-GAGGUGUACGGACCUAUCATT-3’; si*SRSF1*, 5’-AGGACAUUGAGGACGUGUUdTdT-3’ and 5’-AACACGUCCUCAAUGUCCUTT-3’.

### Immunofluorescence

Cells on coverslips were fixed with 4% paraformaldehyde at room temperature (RT) for 10 min. Following fixation, the cells were permeabilized using 0.1% Triton X-100 in PBS, blocked with 3% BSA and then incubated with the following primary antibodies: mouse anti-SRSF1 (Invitrogen; 32-4500), rabbit anti-SFPQ (Proteintech; 15585-1-AP), rabbit anti-FBL (Proteintech; 16021-1-AP), mouse anti-FBL (Proteintech; 66985-1-Ig), mouse anti-UBF (Santa Cruz Biotechnology; SC-13125), mouse anti-NPM1 (Invitrogen; 32-5200), mouse anti-Flag (GNI; GNI4110-FG), rabbit anti-γH2AX (Abclonal; AP0687), mouse anti-rRNA (Santa Cruz Biotechnology; SC-33678), mouse anti-p53 (Proteintech; 60283-1-Ig) and rabbit anti-NCL (Proteintech; 10556-1-AP) at 4°C for overnight. After primary antibody incubation, the cells were incubated with Alexa Fluor-labelled secondary antibody at RT in the dark for 1 hour. Finally, the samples were mounted on slides using Fluoromount-G mounting medium (Southern Biotech; 0100-01). Imaging was performed on a Zeiss LSM-900 confocal microscope equipped with a 63× oil immersion objective.

### Immunoblotting

Cells were collected and lysed with ice cold lysate buffer containing 20 mM Tris-HCl [pH 7.5], 150 mM NaCl, 1 mM EDTA, 1 mM EDTA, 1% Triton X-100, 2.5 mM sodium pyrophosphate, 1 mM β-glycerol phosphate and a protease inhibitor cocktail Then samples were centrifuged at 10,000 rpm for 15 min at 4°C. The collected supernatant was boiled at 100°C for 10 min. The protein samples were loaded in SDS-PAGE gel, and then transferred to polyvinylidene fluoride (PVDF) membranes after electrophoresis. The membranes were blocked with 5% milk and then incubated at 4°C for overnight with the following primary antibodies: mouse anti-Flag (GNI; GNI4110-FG), mouse anti-SRSF1 (Invitrogen; 32-4500), rabbit anti-SFPQ (Proteintech; 15585-1-AP), rabbit anti-FBL (Proteintech; 16021-1-AP), mouse anti-NPM1 (Invitrogen; 32-5200), mouse anti-FUS (Proteintech; 68262-1-Ig), mouse anti NPAT (abcam; ab70595), rabbit anti-U1-70K (abcam; ab83306), rabbit anti-α-Tubulin (CST; 2125S), rabbit anti-NCL (Proteintech; 10556-1-AP), Streptavindin-HRP (APEBIO; K1229), rabbit anti-mCherry (Proteintech; 26765-1-AP), rabbit anti-Vinculin (Proteintech; 26520-1-AP), and rabbit anti-DDX5 (Diagbio; db11221). The membranes were subsequently incubated with Horseradish peroxidase-conjugated Goat Anti-Rabbit (JacksonImmunoResearch; 111-035-003) or Horseradish peroxidase-conjugated Goat Anti-Mouse (JacksonImmunoResearch; 115-035-003). Proteins were detected using Immobilon Western HRP Substrate (Merck Millipore; WBKLS0050).

### Generation of KI cell lines with CRISPR/Cas9

KI cell lines were constructed by CRISPR/Cas technology including U-2 OS cells with endogenous SFPQ, FBL, NPM1, DDX5, TCOF1 and U1-70K tagged by mCherry, and endogenous SRSF1 tagged by GFP, as well as HeLa, 293T, and HCT 116 cells with endogenous SFPQ tagged by mCherry. To facilitate homologous recombination, we used 750 bp flanking homologous arms of the target genes as the amplification templates and engineered them into the pUC19 vector (Addgene; 50005). The guide RNA sequences were 5’-CCACAGACATGTCTCGGGAT-3’ for *SFPQ*; 5’-GTTGTGAACGCCGCGGACTC-3’ for *FBL*; 5’-AGGGTCTATATATAAGCGCG-3’ for *NPM1*; 5’-ATATACTTCTAAAGTCTTAT-3’ for *DDX5*; 5’-GTCATAGCAGATCTCGCTCT-3’ for *SRSF1*; 5’-CCGGCCGG-GGGTCGCGGGTA-3’ for TCOF-1; 5’-GGAGAATGGGTATTTGATGG-3’ for *U1-70K*. Meanwhile, these sgRNA sequences were cloned into the PX459 vector. After transient transfection, the selected pools were directly sorted into 96-well cell culture plates by flow cytometry, and the KI efficiency was determined via targeted PCR and DNA sequencing.

### Proximity labeling by TurboID

Cells stably expressing TurboID fused in frame with target proteins were incubated with 500 μM biotin for 120 min, followed by affinity pull-down using streptavidin-conjugated magnetic beads (GenScript; L00936). The mixture was rotated at 10 rpm for 3 hours at 4°C. Subsequently, the magnetic beads were eluted with a buffer containing 50 mM Tris-HCl (pH = 7.5) and 1% SDS, and boiled at 100°C for 10 min. The elution was analyzed by immunoblotting to confirm pull-down efficiency, while the remaining was subjected to LC-MS/MS analysis (Bruker). Mass spectrometry data were processed using Perseus, and protein expression changes between AMD-treated and untreated samples were compared to identify differentially expressed proteins.

### RNA-sequencing

Total RNA was extracted using the AG RNAex Pro Reagent (Accurate Biology; AG21102). RNA-seq was performed by Tsingke Biotechnology Co., Ltd. (Beijing, China). RNA quality and integrity were assessed using the NanoDrop spectrophotometer and the LabChip GX Touch system (Revvity), respectively. RNA concentration was then quantified with a Qubit 3.0 fluorometer using the Qubit™ RNA Broad-Range Assay Kit (Thermo Fisher; Q10210). To enrich PCR products of 200–500 bp in length, library preparation was carried out using the KC™ Digital mRNA Library Preparation Kit (Seqhealth Technology Co., Ltd., Wuhan, China). After quantification, sequencing was performed on the DNBSEQ-T7 platform (MGI).

For quantitative analysis of the RNA-seq data, adapters and low-quality sequences were removed from the raw reads using fastp (https://github.com/OpenGene/fastp). The filtered reads were then aligned to the Homo sapiens genome (GRCh38.112) using HISAT2 (https://daehwankimlab.github.io/hisat2/). Transcripts were preliminarily assembled and merged using StringTie (https://github.com/gpertea/stringtie). The final assembly annotation was obtained by comparing the assembled transcripts with reference annotations using gffcompare (http://ccb.jhu.edu/software/stringtie/gffcompare.shtml), followed by quantitative analysis. Differential expression analysis between samples was performed with DESeq2 (http://www.bioconductor.org/packages/release/bioc/html/DESeq.html). Alternative splicing events were identified using rMATS (version v3.2.5) with the criteria FDR < 0.05 and |Δψ| ≥ 0.05. Finally, GO enrichment analysis was carried out on all differentially expressed genes associated with alternative splicing.

### FRAP

The Zeiss Cell Discoverer 7 confocal microscope equipped with a water lens was used for FRAP experiments. The target area was pre-bleached for 3 seconds with an argon ion laser (405 nm and 488 nm wavelength) operating at 100% power. Subsequently, the fluorescent regions of interest were selected for photobleaching, and fluorescence measurements were conducted for 150 s or 15 s, with images captured every 7.5 s or 0.75 s, for a total of 20 cycles. The EasyFRAP software (https://easyfrap.vmnet.upatras.gr/) was employed for data normalization and analysis.

### Measurements of pH in cells

To perform ratiometric pH measurements in the DNC, we employed the pH-sensitive fluorescent probe BCECF-AM (Beyotime; S1006). This dye exhibits pH-dependent fluorescence intensity when excited at 490 nm, while remaining stable upon excitation between 400 nm and 440 nm. Intracellular pH was determined by measuring the emission at 535 nm using a Zeiss LSM980 confocal laser scanning microscope (Zeiss) following excitation at 405 nm and 488 nm, respectively. The ratio of fluorescence intensities obtained at these two excitation wavelengths (488 nm/405 nm) was used to calculate pH values.

A standard curve was first generated using 2 μM BCECF-AM in calibrated serum-free McCoy’s 5A medium at pH 5.80, 6.29, 6.88, 7.44, 8.14, and 8.73, respectively. 10 μM Nigericin (Shanghai Yuanye Bio-Technology; S25116) was added to each calibration buffer 5 minutes prior to measurement. The fluorescence intensity ratio (488 nm/405 nm) was plotted against pH to establish the calibration curve.

For nucleolar pH measurements, U-2 OS cells expressing endogenously tagged fluorescent markers were used. Target regions were identified and masked based on the fluorescence signal of these markers. The emission ratio (∼490 nm/∼440 nm) within the nucleolar masks was quantified for each cell. Nucleolar and the DNC pH for each cell were derived by fitting the ratiometric values to the standard curve. Data from each experimental condition were pooled from 2-3 independent biological replicates.

### FISH

FISH was performed according to previously described method^41^. The RNA probes of 5’ ETS used as previously reported^18^: 5’-CTCTCAGATCGCTAGAGAAGGCTTTTCTCACCGAGGGTGGGTCACACTCC-3’. Briefly, cells were washed with PBS, fixed with 4% PFA for 15 min, and permeabilized with 70% ethanol overnight at 4°C. After washed with the wash buffer (2x SSC, 50% formamide) for 5 min at RT, cells were incubated at 37°C overnight in the hybridization solution (2x SSC, 200nMvanadyl-ribonucleoside complex, 50% formamide, 0.02% BSA), containing 10 nM of the RNA FISH probes. The coverslips were washed with the wash buffer at 37°C for 30 min. The coverslips were mounted with mounting medium.

### Quantitative real-time PCR

Total RNA from U-2 OS cells was extracted by AG RNAex Pro Reagent (Accurate Biotechnology; AG21102) according to the manufacturer’s instructions. 1 µg of total RNA was preformed to reverse transcribed by HiScript IV All-in-One Ultra RT SuperMix (Vazyme; R433-01). The product was diluted and mixed with specific primers and 2×Universal SYBR Green Fast qPCR mix (Abclonal; RM21203). We then performed PCR reactions by a Bio-Rad CFX96 real-time PCR system and collected data by Bio-Rad CFX Manager. GAPDH mRNA expression level was served as internal references for normalization, and the ΔΔCt method was applied to determine the abundance of the target genes.

### Measurements of ROS

Intracellular ROS levels were measured by live cell imaging using the redox-sensitive fluorescent probe CM-H_2_DCFDA (Invitrogen; C6827) or the hydrogen peroxide-selective fluorescent probe Pentafluorobenzenesulfonyl fluorescein (Amgicam; ajci73430). Briefly, a 2 μM working solution of CM-H_2_DCFDA or a 2.5 μM solution of Pentafluorobenzenesulfonyl fluorescein was prepared in serum-free culture medium.

Before staining, cells were gently washed with serum-free medium to remove residual compounds. The diluted probe was then directly applied to the cells, which were subsequently incubated in the dark at 37°C for 30 min. Following incubation, cells were washed once or twice with serum-free medium to remove excess extracellular dye. Fluorescent imaging was carried out on a ZEISS LSM-980 confocal microscope with a 63× oil-immersion objective. All images were acquired under standardized exposure settings to ensure consistency across samples.

roGFP1 is a redox-sensitive green fluorescent protein used to detect intracellular H_2_O_2_ levels^29^. To confer DNC-targeting capability, the RRM domain of SRSF1 was fused in frame to roGFP1, ensuring accurate detection of the redox status within the DNCs. Fluorescence imaging was performed using a ZEISS LSM 980 confocal microscope equipped with a 63× oil-immersion objective. mCherry-SFPQ KI U-2 OS cells expressing RRM_SRSF1_-roGFP1 were imaged with excitation at 405 nm and 488 nm, and the corresponding fluorescence signals were collected using bandpass filters of 405-520 nm and 488-520 nm, respectively. All images were acquired under standardized exposure settings to ensure consistency across samples. Quantitative analysis of the 405/488 nm fluorescence ratio was performed using ImageJ.

### Measurements of electrical potential

To measure the DNC potential, the potential-responsive dye DiOC_2_(3) (Maokangbio; MX4008-100MG) was used in live-cell imaging, where its fluorescence intensity is directly proportional to the potential. A 4 mM stock solution of DiOC_2_(3) was prepared in DMSO and then diluted with serum-free McCoy’s 5A medium to a final working concentration of 2 µM. Before imaging, mCherry-SFPQ KI U-2 OS cells were incubated with the dye working solution for 30 min in an incubator at 37°C and 5% CO_2_. The dye solution was then removed and replaced with complete culture medium. Fluorescence imaging was performed using a ZEISS LSM 980 confocal microscope with a 63× oil-immersion objective. The dye requires a pre-activation step: the cells to be imaged are irradiated with a 488 nm laser for 5 seconds. During imaging, a 561 nm excitation light source was used to visualize mCherry-SFPQ expression and thus determine the focal plane of the DNCs. The potential was determined by measuring the emission at 535 nm following excitation at 488 nm, using the same confocal microscope. All images were acquired under standardized exposure settings to ensure consistency across samples. ImageJ was used for quantitative analysis.

### Protein sequence analysis

The physicochemical and functional properties of putative DNC proteome (referred to as Target Group) against the human reference proteome (referred to as Total Group) was analyzed using basic protein data sourced from the UniProt database. A custom Python 3.9 bioinformatics pipeline, employing Biopython, computed six key metrics for each protein: isoelectric point, net charge and net charge per residue at pH 7.4, the proportions of basic amino acids (lysine and arginine), cysteine, and methionine. Systematic comparison between the groups involved a three-step statistical analysis: first, determining the percentile rank of the Target Group’s median within the Total Group’s distribution; second, applying the two-tailed Mann–Whitney U test (α = 0.05) to compare metric distributions for statistical significance; and third, quantifying the practical magnitude of differences using Cliff’s δ effect size. All data manipulation was handled by Pandas, with statistical computations by using Scipy.

### Sucrose gradient ultracentrifugation

We performed sucrose gradient ultracentrifugation as previously described^42^. Briefly, a 5-50% continuous sucrose gradient was prepared in a 14 mL ultracentrifuge tube (Beckman Coulter) by sequentially layering sucrose solutions from 50% to 5% in 5% decrements. Each step consisted of 1.2 mL of sucrose solution prepared in gradient buffer (10 mM Tris pH 7.4, 100 mM NaCl, 1 mM EDTA), with each layer frozen at -80°C for 15 min before addition of the next.

U-2 OS cells grown on 15 cm dishes were lysed, and protein concentration was quantified. To remove RNA, equal amounts of protein samples (∼500 mg) were digested on ice for 1 hour using a cocktail of RNases, including 1 mg of RNase A (LABLEAD; R4875), 5 U of RNase I (Thermo Fisher; EN0601), 500 U of RNase T1 (Beyotime; R7096S), and 1 U of RNase III (Beyotime; R7086S). After digestion, samples were loaded onto pre-chilled 5-50% sucrose gradients and subjected to ultracentrifugation in a Beckman rotor at 40,000× g for 18 hours at 4°C.

Following centrifugation, 12 equal-volume fractions (1.2 mL each) were collected from the top of each gradient. Proteins in each fraction were precipitated by adding 0.6 mL of absolute ethanol and incubating at -20°C overnight. Precipitated proteins were pelleted by centrifugation at 13,000× g for 30 min at 4°C, washed once with 70% ethanol, and air-dried. Pellets were resuspended in 2× SDS sample buffer, boiled at 95°C for 10 min, and equal volumes from each fraction were analyzed by SDS-PAGE and immunoblotting with the indicated antibodies.

### Measuring the DNC polarity in cells

To establish a calibration curve for fluorescence lifetime (*τ*_AvInt_) as a function of the dielectric constant (ε), we incorporated SBD into mixtures of methanol and 1,4-dioxane with known ε values and measured the corresponding *τ*_AvInt_ of SBD. Data points for *τ*_AvInt_ were obtained from 11 groups of SBD in solvents of varying dielectric constants, and a standard curve was generated by fitting these values to the respective dielectric constants.

For FLIM experiments, cells were transfected with 1 μg of plasmids encoding HaloTag-fused SRSF1 or SFPQ to label the DNCs, or HaloTag-fused TCOF1 to label FC. When cells reached approximately 40% confluency, half of the culture medium (DMEM) was replaced with fresh medium supplemented with SBD-Halo. After 24 hours of incubation, cells were gently washed with Gibco™ DMEM/F-12, no phenol red (Gibco; 21-041-025) containing 10% FBS to remove unbound HaloTag substrate and reduce background fluorescence, followed by treatment with 1 μM AMD for 4 hours. Live-cell FLIM was carried out on a Leica STELLARIS 8 FALCON confocal microscope equipped with a pulsed white-light laser (WLL) operating at 448 nm excitation and a 10 MHz repetition rate. During imaging, cells were maintained under physiological conditions (37°C, 5% CO_2_) using a climate-controlled chamber. Regions of interest (ROIs) were defined using the LAS X FLIM/FCS software, and fluorescence lifetime histograms and decay curves were generated. The fluorescence decay data were fitted using a multi-exponential reconvolution model, and the intensity-weighted mean fluorescence lifetime (*τ*_AvInt_) was extracted as a quantitative indicator of local polarity.

### Fluorescence correlation spectroscopy

FCS measurements were carried out at 20°C using a MicroTime 200 system (PicoQuant, Germany) operated in FCS mode. Cells were seeded on glass-bottom culture dishes (Cellvis, USA) and treated with AMD + DMSO or AMD + NAC for 4 h prior to measurements. A 532 nm pulsed laser (40 MHz) set at 15 µW was used for excitation. The laser beam was focused 3-5 µm above the coverslip to probe droplets within the nuclear compartment, through a 60×/1.20 W water-immersion objective (UPlanApo, Olympus, Japan). Emitted fluorescence was filtered with a 620/60 nm bandpass filter (Chroma, USA), and detected with single-photon avalanche diodes. Correlation curves were generated with SymPhoTime64 software v2.7 (PicoQuant, Germany) and fitted using the 3D triplet-extended model to extract diffusion times.

### FCS data analysis

In FCS, the normalized autocorrelation function (ACF) is defined by

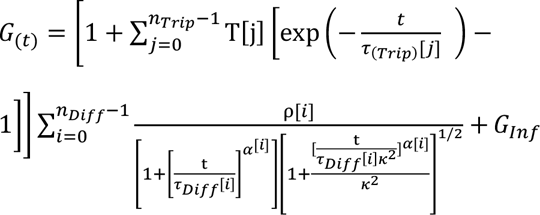

Where T[j] is dark (triplet) fraction of molecules, *τ*_(_*_Trip_*)[*j*] is lifetime of the dark (triplet) states, *n_Trip_* is number of dark (triplet) states, *n_Diff_* is number of independently diffusing species. ρ[*i*] is the contribution of the *i^t^*^ℎ^ diffusing species. *τ_Diff_*[*i*] is the diffusion time of the *i^t^*^ℎ^ diffusing species. α is anomaly parameter of the i^th^ diffusing species. *κ* is the length to diameter ratio of the focal volume. *G_Inf_* is correlation offset.

Diffusion constant of the k^th^ diffusing species (D), average number of molecules in the focal volume (N) and concentration of molecules in the focal volume (C) were corrected in this work using the calculated factors and the following equation:

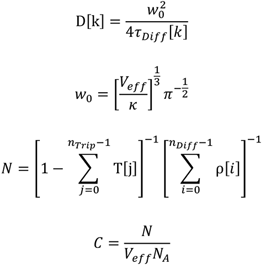

Where w_0_ is effective lateral focal radius at ^1^/e^2^ intensity, *V*[*eff*] is effective excitation volume, *N_A_* is the Avogadro constant.

### Image process and quantitative analysis

The partition coefficient, defined as the ratio of protein concentration in the dense phase to that in the dilute phase, was quantified for protein pairs using the Coloc2 plugin in ImageJ. Whole-cell measurements were performed on images pre-processed by conversion to 8-bit format, followed by Gaussian filtering. To compute this coefficient, a three-mask strategy was employed: (1) a dense-phase mask (ROI1) generated from the specified channel; (2) a corresponding mask for the target protein signal (ROI2) from its respective channel, used to obtain the mean intensity within the dense phase; and (3) a derived dilute-phase mask (ROI3), whose mean intensity was calculated from the total intensity and area data of ROI2. The final partition coefficient was then calculated as (Mean Intensity of ROI2)/(Mean Intensity of ROI3).

Pearson’s correlation is defined as the degree of linear correlation between the fluorescence signals of two different channels within the same focal plane. Analysis was performed using the JaCoP plugin in ImageJ, following a standardized pre-processing workflow: images from both channels were first converted to 8-bit format, then a Gaussian filter was applied to reduce background noise, and finally quantitative analysis was carried out using Pearson’s coefficient component within the JaCoP plugin.

Fluorescence intensity measurements were performed across the entire cell on the focal plane using the Coloc2 plugin in ImageJ. The analysis followed a standardized pre-processing workflow: images were first converted to 8-bit format, then a Gaussian filter was applied to reduce background noise, and finally, a threshold was set based on signal intensity to define the regions for quantification.

Nucleolar size measurements were performed by manual image segmentation using ImageJ software. Briefly, images were converted to 8-bit format, followed by Gaussian filtering to reduce background noise, and thresholding was applied based on signal intensity. Detected objects in each image were then manually verified, and only nucleoli located in the focal plane-identified by a thin, low-contrast boundary-were selected for subsequent size analysis.

## Quantification and statistical analysis

All analysis was performed in a blinded manner with respect to the experimental conditions. Unless otherwise specified, data in this study are expressed as mean ± SD with data pooled from at least two independent biological replicates, or as mean ± SEM from at least three independent biological replicates. Quantifications-including those of partition coefficient, Pearson’s correlation, pH, and fluorescence intensity-were performed on a cell-by-cell basis. Statistical significance was evaluated using two-tailed unpaired Student’s *t*-test or ANOVA, as appropriate, with *p < 0.05 considered statistically significant. Data analysis was carried out using GraphPad Prism 10 software. Representative microscopy and western blot images shown were derived from at least two independent experiments. Detailed statistical parameters are provided in the figures and accompanying legends.

## Supplementary information

**Figure S1. Related to Figure 1.**
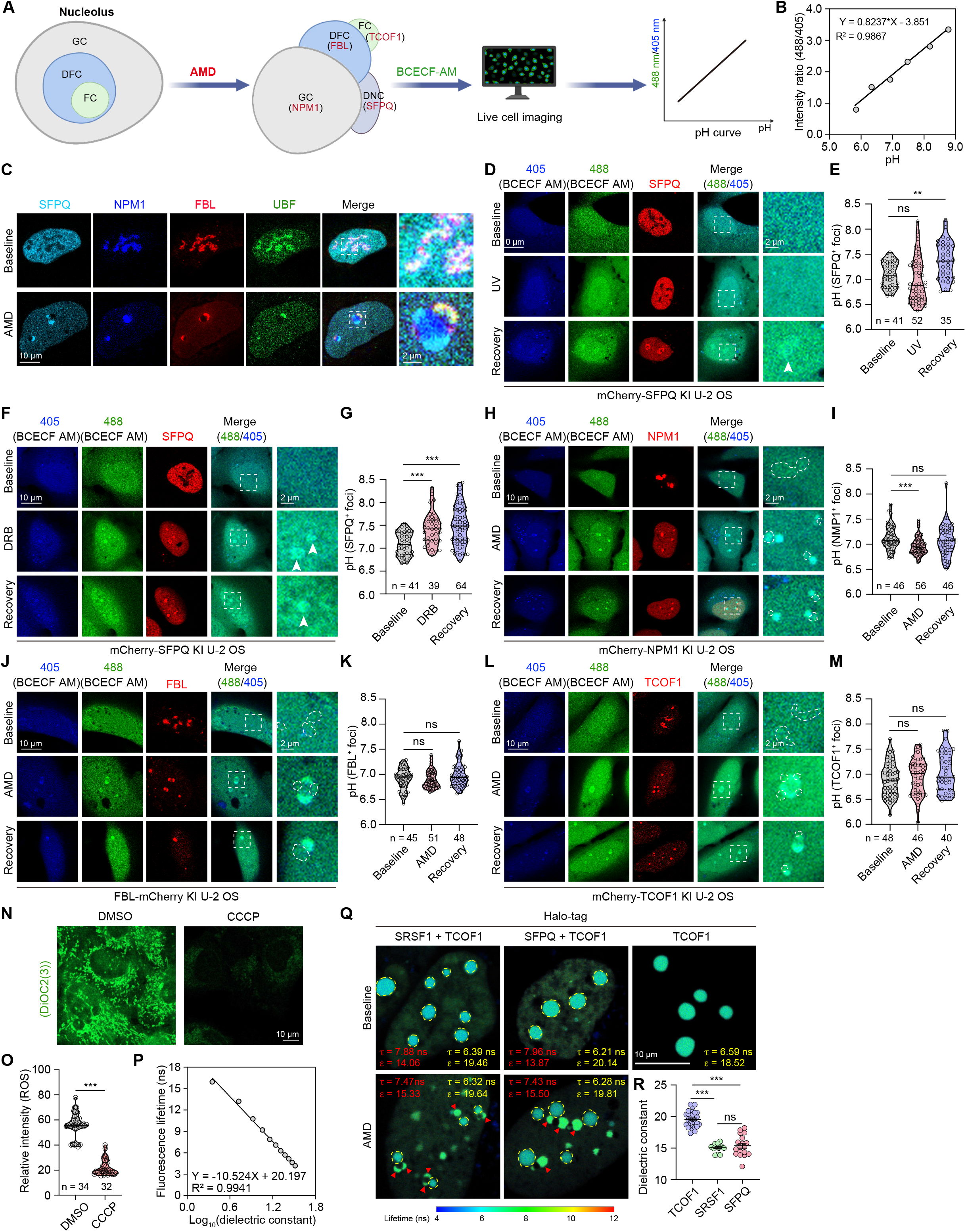
DNCs enable redox reactions and produces ROS by establishing interfacial pH gradient. (**A**) Schematic illustration of pH measurement using the fluorescent dye BCECF-AM. **(B)** Standard curve for pH measurement in cells by pH-sensitive BCECF-AM dye. Linear progression was established between the pH values of standard buffers and the ratiometric intensity of 488/405 nm. **(C)** Localization of SFPQ, NPM1, FBL and UBF in U-2 OS cells 4 hours after treatment with AMD. (**D-G**) Live-cell imaging was performed for mCherry-SFPQ KI U-2 OS treated with UV (**D**, **E**) or DRB (**F**, **G**), then recovered under normal growth conditions with BCECF-AM. ns, not significant; **p < 0.01; ***p < 0.001 by one-way ANOVA (Kruskal-Wallis test). (**H-M**) Live-cell imaging was performed for NPM1-mCherry (**H**, **I**), FBL-mCherry (**J**, **K**), or mCherry-TCOF (**L**, **M**) KI cells with the pH indicator BCECF-AM. These U-2 OS cells were first treated with AMD, and then recovered under normal growth conditions. pH was quantified by calculating the fluorescence intensity ratio at 488/405 nm wavelengths. Dotted circles denote areas positive for mCherry signal. ns, not significant; *p < 0.05; **p < 0.01; ***p < 0.001 by one-way ANOVA (Kruskal-Wallis test). **(N, O)** Live-cell imaging was performed on DMSO-or CCCP-treated U-2 OS cells by using DiOC_2_(3). The fluorescence intensity of DiOC_2_(3) was quantified (**O**). ***p < 0.001 by unpaired Student’s *t*-test. (**P**) The calibration curve for fluorescence lifetime as a function of the dielectric constant. **(Q, R)** FLIM images for SBD-labeled SRSF1, SFPQ and TCOF-1 subcompartments. Polarity was quantified (**P**). ns, not significant; ***p < 0.001 by one-way ANOVA.

**Figure S2. Related to Figure 1.**
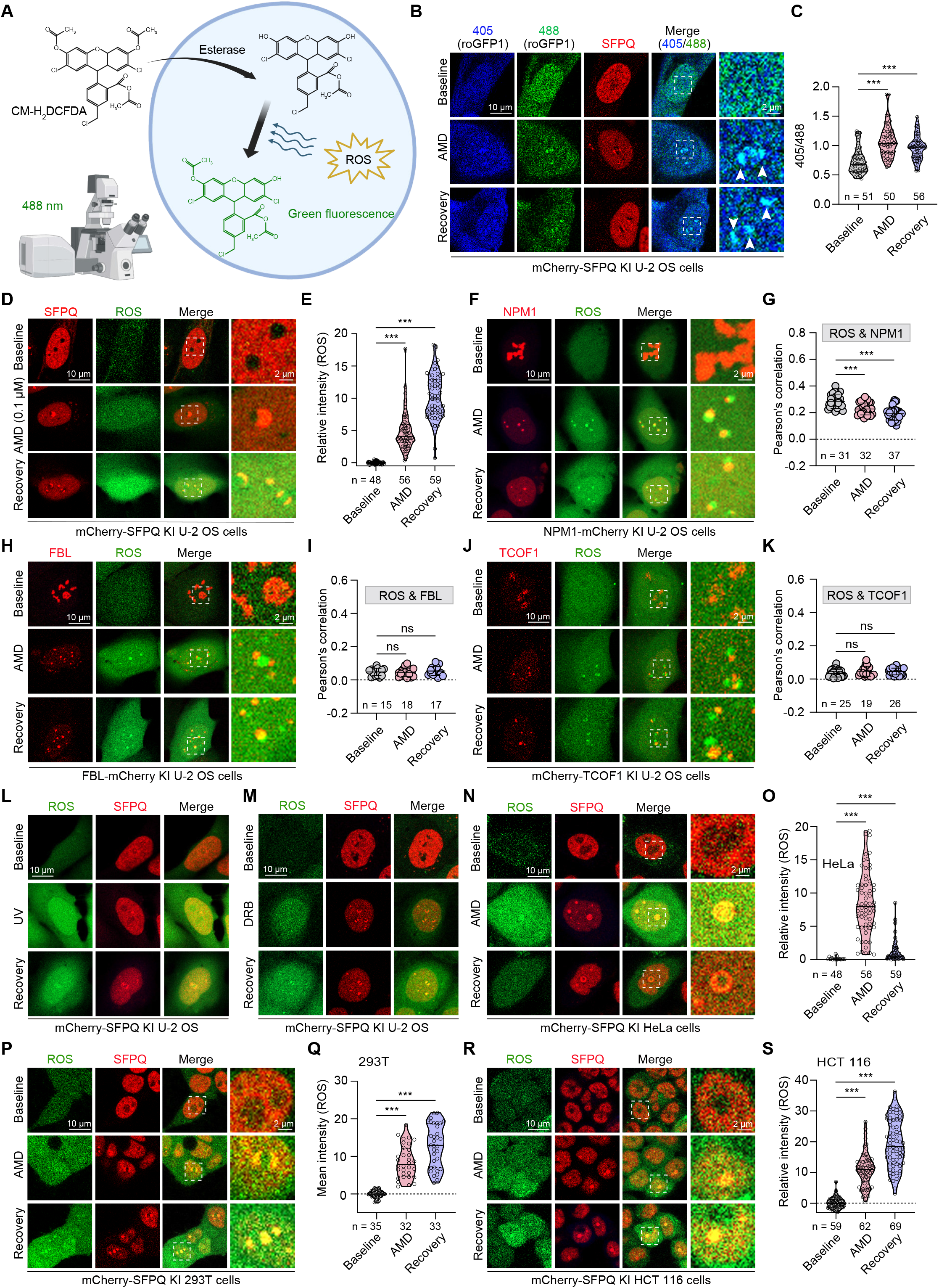
DNCs enable redox reactions and produces ROS by establishing interfacial pH gradient. (**A**) Schematic illustration of ROS measurement using the fluorescent dye CM-H_2_DCFDA. **(B, C)** Live cell imaging of mCherry-SFPQ KI U-2 OS expressing RRM_SRSF1_-roGFP1 (for DNC targeting) treated with AMD and recovered in normal growth condition. ROS was quantified by calculating the ratio of the fluorescence intensity of 405/488 nm in the DNCs marked by mCherry-SFPQ (**C**). **(D, E)** Live-cell imaging was performed on mCherry-SFPQ KI U-2 OS cells following 0.1 µM AMD treatment and subsequent recovery under normal growth conditions, utilizing the ROS-sensitive probe CM-H_2_DCFDA. The fluorescence intensity of ROS was quantified (**E**). (**F-K**) Live-cell imaging was conducted in U-2 OS cells expressing NPM1-mCherry (**F**, **G**), FBL-mCherry (**H, I**), or mCherry-TCOF1 KI (**J, K**). Cells were treated with AMD and returned to normal growth conditions and incubated with the ROS indicator CM-H_2_DCFDA. Pearson’s correlation of the ROS signal and the different nucleolar subphases was quantified in (**G**), (**I**), and (**K**), respectively. (**L, M**) Live-cell imaging of mCherry-SFPQ KI U-2 OS cells treated with UV (**L**) or DRB (**M**), followed by recovery under normal growth conditions in the presence of CM-H_2_DCFDA. (**N–S**) Live-cell imaging of mCherry-SFPQ KI HeLa (**N, O**), 293T (**P, Q**), or HCT 116 (**R, S**) cells treated with AMD followed by recovery under normal growth conditions with CM-H_2_DCFDA. For (**C**), (**E**), (**G**), (**I**), (**K**), (**O**), (**Q**), and (**S**), ns, not significant; *p < 0.05; **p < 0.01; ***p < 0.001 by one-way ANOVA.

**Figure S3. Related to Figure 2.**
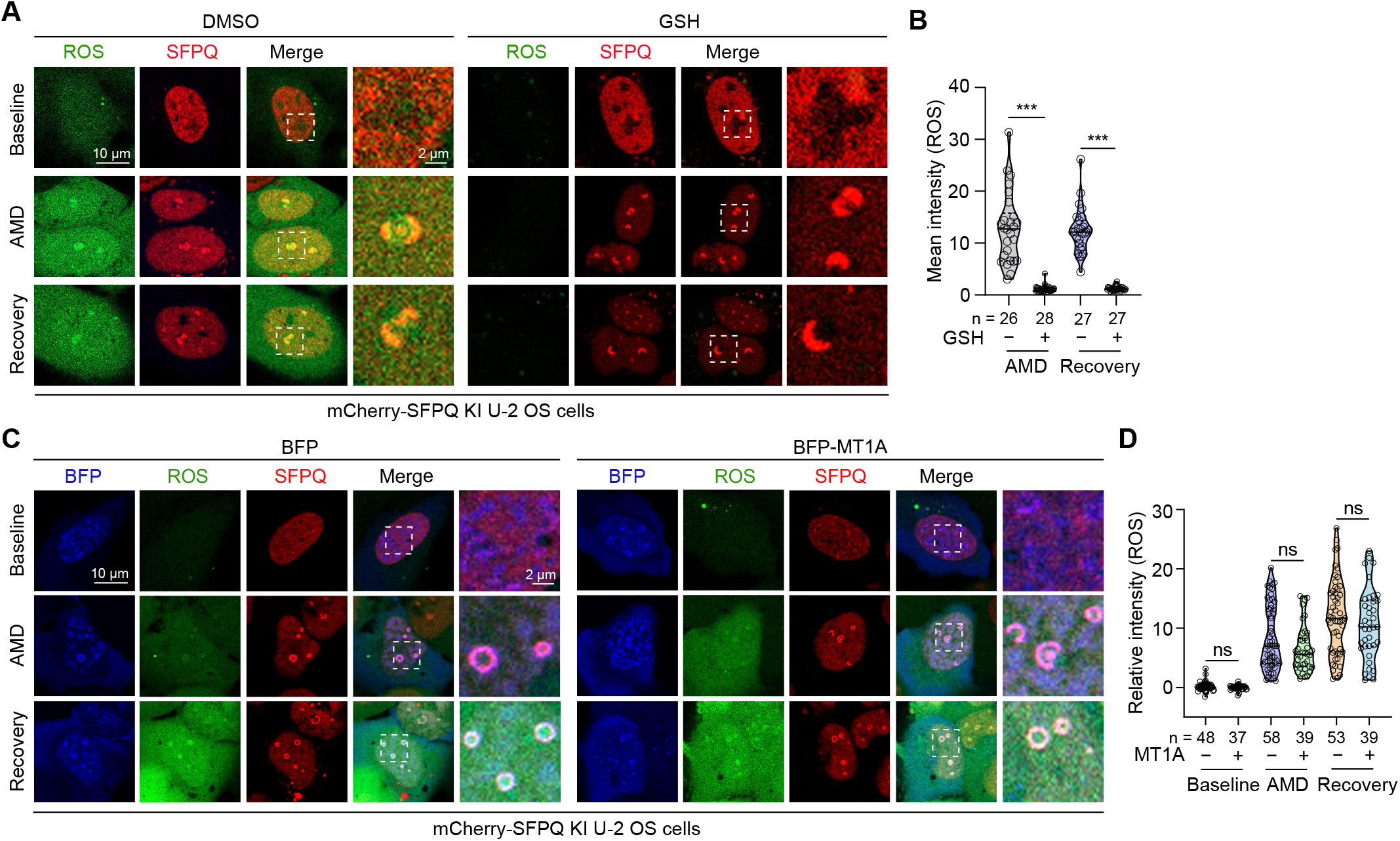
Uncovering the chemical mechanisms of ROS generation in the DNCs. (**A, B**) Live-cell imaging of mCherry-SFPQ KI U-2 OS cells treated with AMD in the presence of DMSO or NAC, and subsequently recovered under normal growth conditions. The fluorescence intensity of ROS was quantified (**B**). (**C, D**) Live-cell imaging was performed on mCherry-SFPQ KI U-2 OS cells expressing BFP-MT1A following AMD treatment and subsequent recovery under normal growth conditions. The fluorescence intensity of ROS was quantified (**D**). For (**B**) and (**D**), statistical significance was determined using a two-tailed unpaired Student’s *t*-test. ns, not significant; ***p < 0.001.

**Figure S4. Related to Figure 3.**
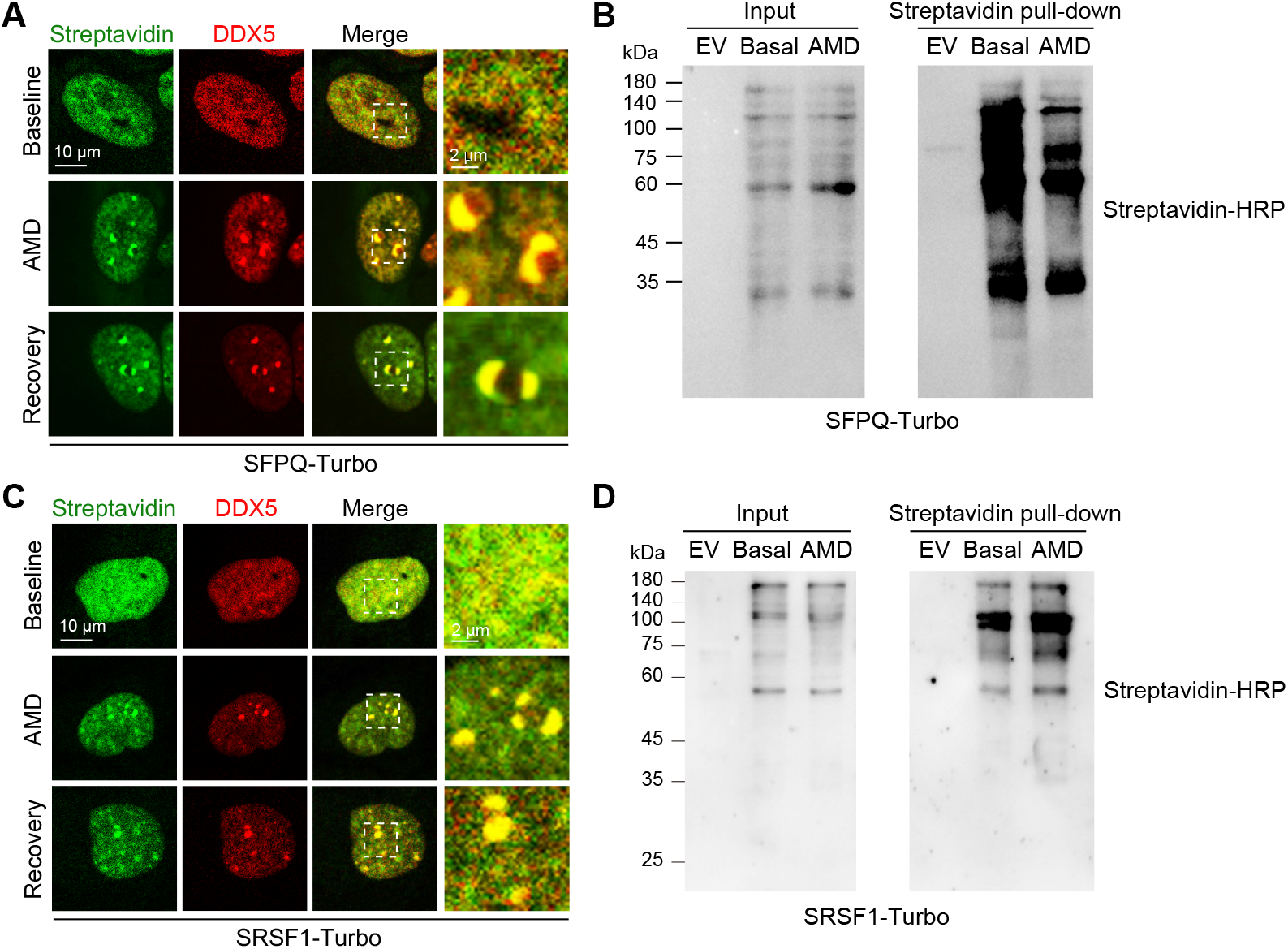
DNC proteome is enriched with positively charged residues. (**A**) WT U2-OS cells stably expressing SFPQ-Turbo were treated with AMD and then recovered under normal growth conditions. Subsequently, cells were subjected to fluorescence staining with YSFluor™ 647-conjugated Streptavidin. **(B)** U2-OS cells transfected with SFPQ-Turbo were subjected to affinity capture with streptavidin magnetic beads, followed by immunoblotting analysis. **(C)** U2-OS WT cells expressing SRSF1-Turbo were treated with AMD, allowed to recover under normal growth conditions, and then labeled by Fluor™ 647-conjugated Streptavidin. **(D)** U2-OS cells transfected with SRSF1-Turbo were subjected to affinity capture with streptavidin magnetic beads, followed by immunoblotting analysis.

**Figure S5. Related to Figure 4.**
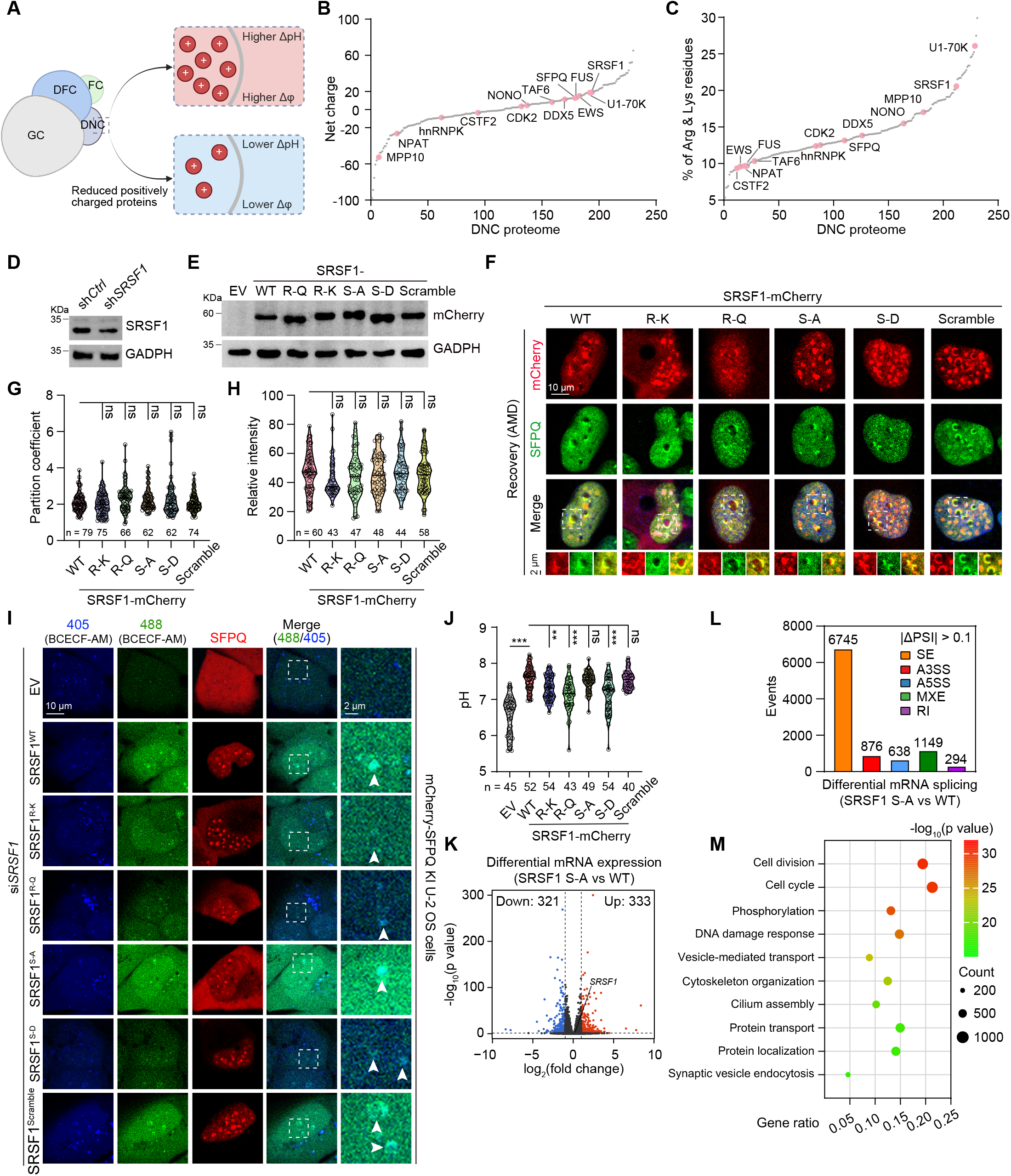
SRSF1 promotes ROS production through tuning the interfacial pH gradient in the DNCs. (**A**) Schematic diagram illustrating the proposed hypothesis. (**B, C**) Bioinformatics analyses of the total net charge (**B**) and proportion of arginine and lysine residues of the putative DNC proteome (**C**). (**D**) Immunoblot analysis of lysates prepared U-2 OS cells transduced with sh*Ctrl* or sh*SRSF1*. (**E**) Immunoblot analysis of lysates prepared from 293T cells expressing EV (empty vector), WT (wild-type) SRSF1, or SRSF1 mutants. (**F, G**) U2-OS WT cells expressing the indicated SRSF1 mutants were treated with AMD and recovered under normal growth conditions. Cells were further processed for immunostaining using the indicated antibodies (**F**). Partition coefficient of SRSF1 mutants in the DNCs was quantified (**G**). (**H**) The relative fluorescence intensity of SRSF1 mutants in the DNCs was quantified from live cell experiments. (**I, J**) WT U-2 OS cells transduced with si*SRSF1* were reconstituted with siRNA-resistant SRSF1 variants. These cells were treated with AMD and allowed to recover for 2 hours, after which the DNC pH was measured by live cell imaging. (**K**) The volcano plot shows the differential mRNA expression between U-2 OS cells expressing SRSF1 WT and S-A. (**L**) Differential alternative splicing events of mRNA between U-2 OS cells expressing SRSF1 WT and S-A. Events with |ΔPSI (percentage spliced in)| > 0.1 are shown. A3SS, alternative splice site; A5SS, alternative 50 splice site; MXE, mutually exclusive exon; RI, retained intron; SE, skipped exon. (**M**) Gene ontology (GO) analysis of differential alternative splicing events of mRNA between U-2 OS cells expressing SRSF1 WT and S-A. For (G), (H), and (J), statistical significance was determined using one-way ANOVA, followed by Dunnett’s test. ns, not significant; *p < 0.05; **p < 0.01; ***p < 0.001.

**Figure S6. Related to Figure 5.**
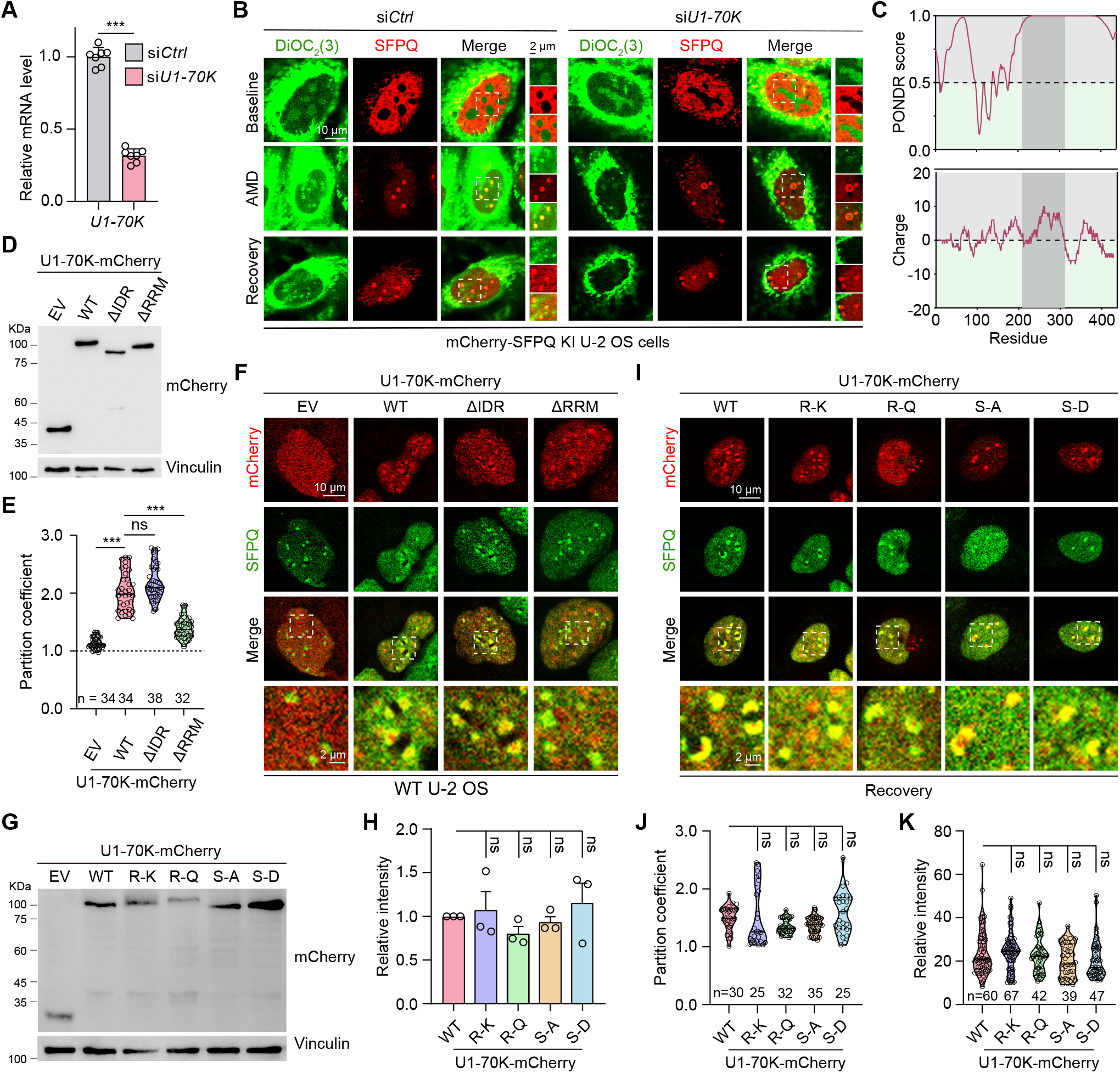
U1-70K is required for ROS synthesis through establishing the interfacial pH gradient in the DNCs. (**A**) WT U-2 OS cells were transfected with si*Ctrl* or si*U1-70K*. *U1-70K* mRNA expression level was determined by qRT-PCR. Data are shown as mean ± SD. n = 7 technical replicates. (**B**) Live-cell imaging of mCherry-SFPQ KI U-2 OS cells transfected with siCtrl or siU1-70K, treated with AMD, and recovered under normal growth conditions in the presence of DiOC_2_(3). (**C**) Predictor of natural disordered regions (PONDR) and charge plot of U1-70K. (**D**) 293T cells transfected with the indicated constructs were subjected to Immunoblotting analysis. (**E, F**) U2-OS WT cells expressing the indicated U1-70K truncations were treated with AMD and recovered under normal growth conditions. Cells were further processed for immunostaining using the indicated antibodies (**F**). Partition coefficient of U1-70K truncations in the DNCs was quantified (**E**). (**G, H**) 293T cells transfected with the indicated constructs were subjected to Immunoblotting analysis. Data are shown as mean **±** SEM. n = 3 biological replicates. (**I, J**) U2-OS WT cells expressing the indicated U1-70K mutants were treated with AMD and recovered under normal growth conditions. Cells were further processed for immunostaining using the indicated antibodies. Partition coefficient of U1-70K mutants in the DNCs was quantified (**J**). (**K**) Fluorescence intensity of U1-70K mutants in the DNCs was quantified in live cell experiments. For (**A**), statistical significance was determined using a two-tailed unpaired Student’s *t*-test. For (**E**), (**H**), (**J**), and (**K**), statistical significance was determined using one-way ANOVA. ns, not significant; ***p < 0.001.

**Figure S7. Related to Figure 6.**
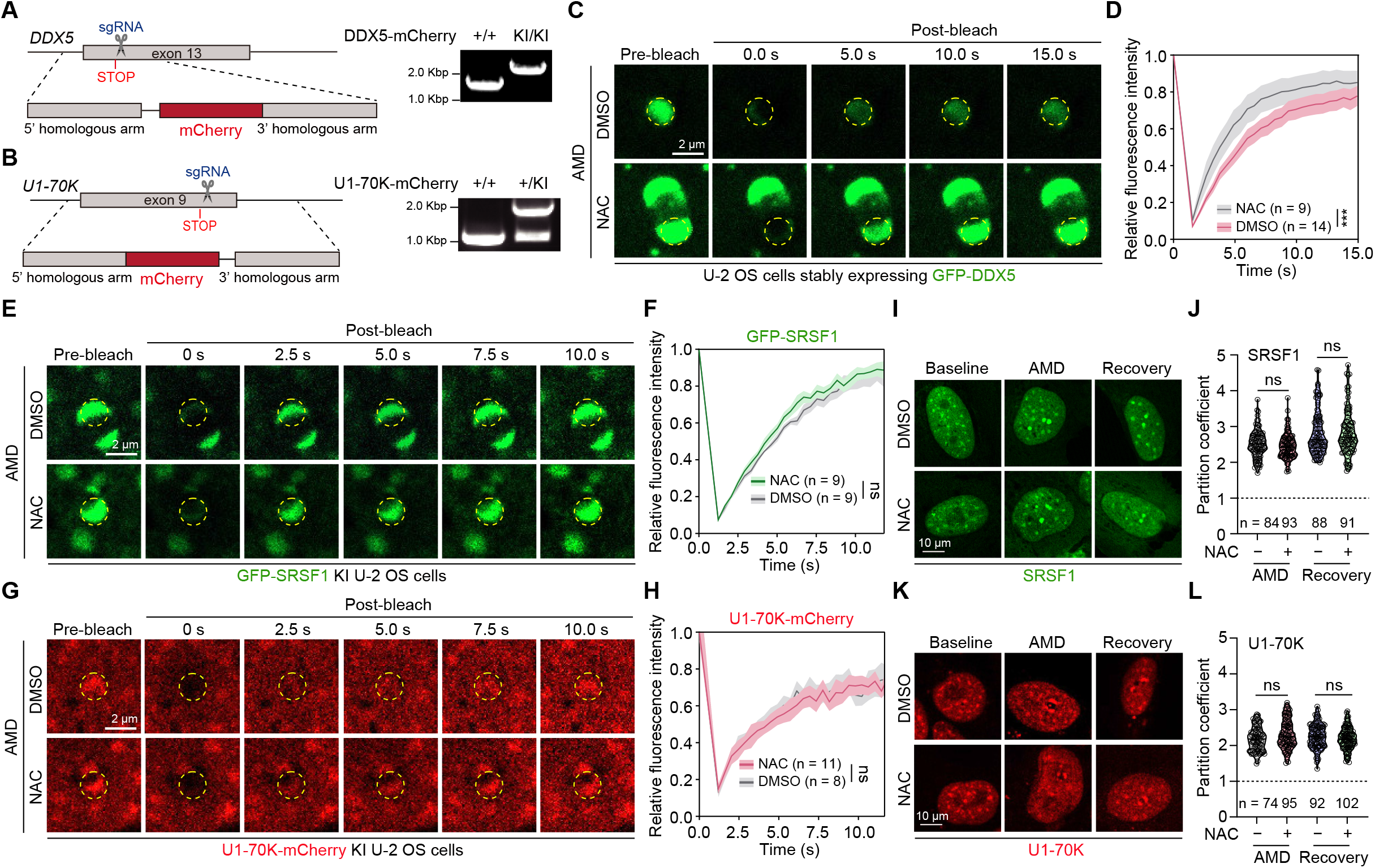
DNC-derived ROS modulates the biophysical properties of redox-reactive DNC constituents. (**A, B**) Schematic illustration and PCR screening of DDX5-mcherry KI **(A)** and U1-70K-mcherry KI **(B)**. (**C**-**H**) U-2 OS cells from DMSO and NAC-treated conditions were subjected to a 4-hour AMD treatment, followed by a 2-hour recovery. FRAP was then performed to monitor and quantitatively analyze the time-dependent changes in fluorescence intensity of GFP-DDX5 (**C**, **D**), GFP-SRSF1 (**E**, **F**), or U1-70K-mCherry (**G**, **H**). Data are presented as mean ± SD. ns, not significant; ***p < 0.001 by two-way ANOVA. (**I**–**L**) U-2 OS cells from DMSO and NAC-treated conditions were treated with AMD for 4 hours and allowed to recover for 2 hours. The partition coefficient was quantified for SRSF1 (**I**, **J**) or U1-70K (**K**, **L**). ns, not significant; *p < 0.05 by a two-tailed unpaired Student’s *t*-test.

**Figure S8. Related to Figure 6.**
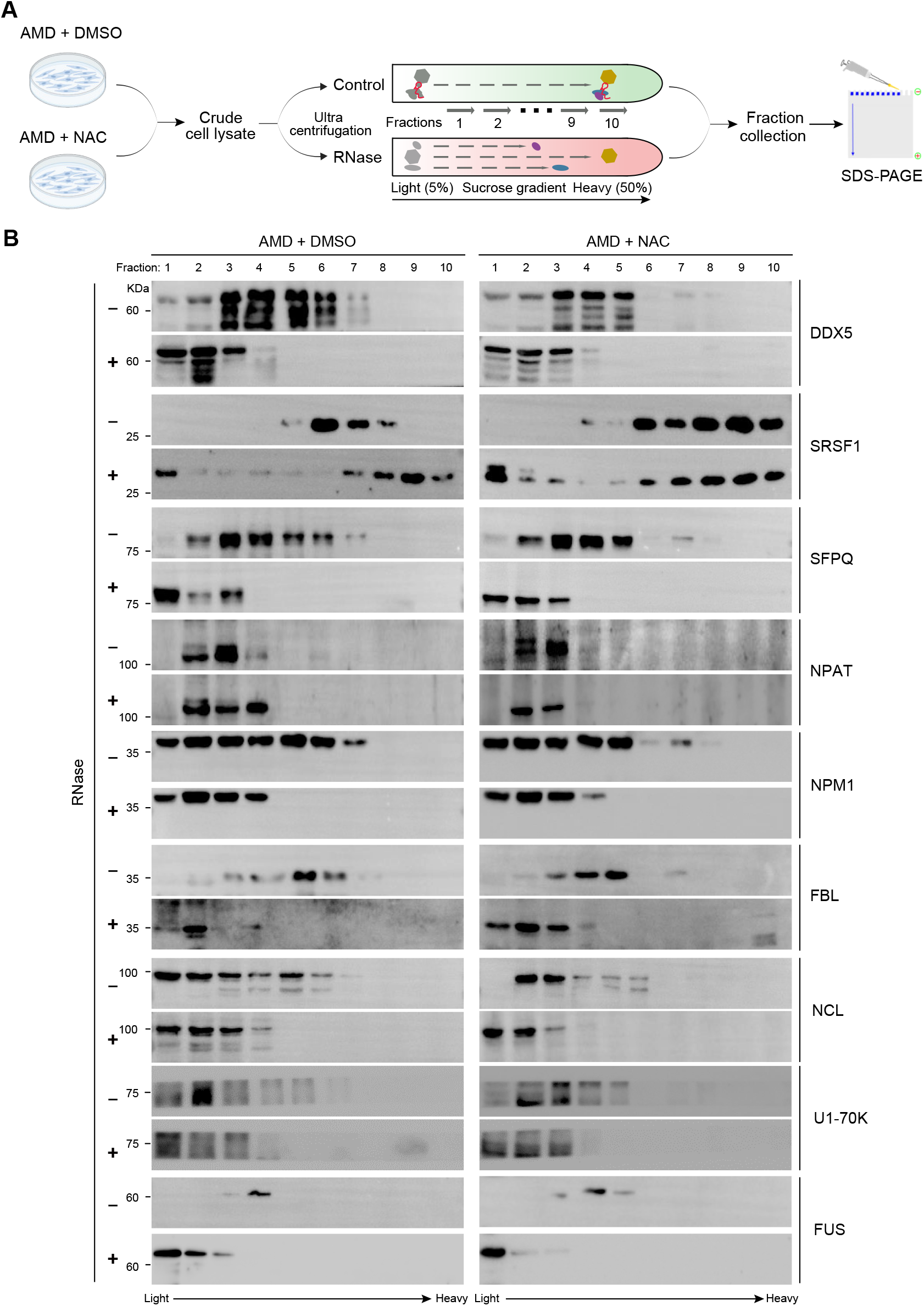
DNC-derived ROS modulates the biophysical properties of redox-reactive DNC constituents. (**A**) Schematic illustration of the experimental design. (**B**) Immunoblot analysis of nucleolar (NPM1, FBL, NCL) and DNC (DDX5, SRSF1, SFPQ, NPAT, U1-70K, FUS) proteins from sucrose gradient fractions after ultracentrifugation with or without RNase treatment in U-2 OS cells under AMD + DMSO and AMD + NAC conditions.

**Figure S9. Related to Figure 7.**
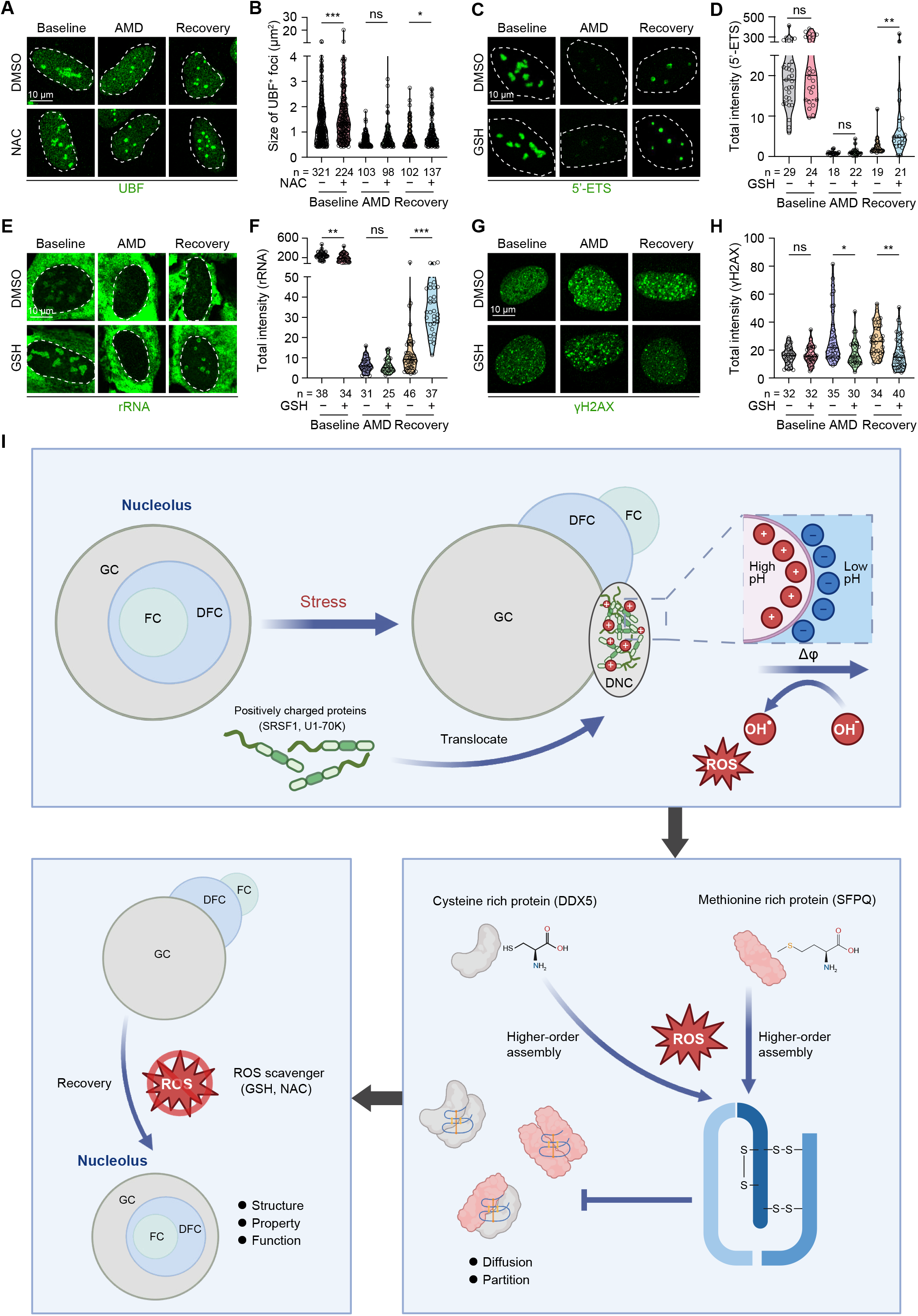
Eliminating ROS accelerates the structural and functional restoration of the nucleolus. (**A, B**) DMSO and NAC-treated U-2 OS cells were exposed to AMD for 4 hours, followed by a 24-hour recovery period. Cell were subjected to immunostaining using an antibody against UBF. The size of UBF foci were then quantified (**B**). DMSO and GSH-treated U-2 OS cells were exposed to AMD for 4 hours and recovered for 24 hours. The cells were subjected to either FISH using a probe for 5’-ETS (**C, D**), immunostaining using antibodies against rRNA (**E, F**), or γH2AX (**G, H**), respectively. The total intensity of 5’-ETS (**D**), rRNA (**F**), and γH2AX (**H**) were quantified, respectively. (**I**) Proposed model of this study. For (**B**), (**D**), (**F**) and (**H**), statistical significance was determined using a two-tailed unpaired Student’s *t*-test. ns, not significant; *p < 0.05; **p < 0.01; ***p < 0.001.

**Table S1.**
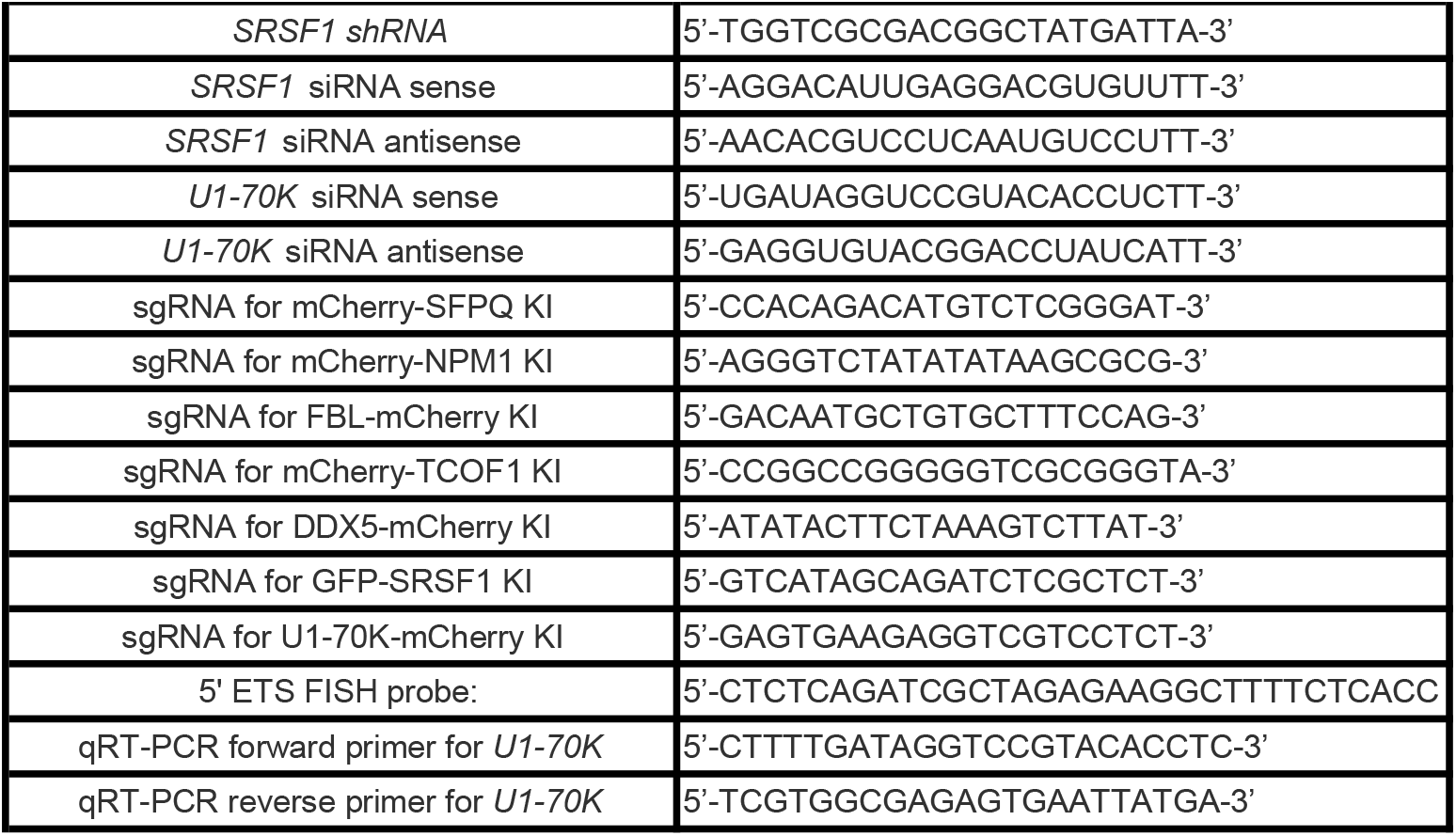
Oligonucleotides used in this study, related to STAR Methods.

